# Influenza A virus polymerase co-opts distinct sets of host proteins for RNA transcription or replication

**DOI:** 10.1101/2025.06.06.658254

**Authors:** Amalie B. Rasmussen, Olivia C. Swann, Ksenia Sukhova, Nancy Liu, Maryn D. Brown, Laura Martin-Sancho, Carol M. Sheppard, Wendy S. Barclay

**Affiliations:** Department of Infectious Disease, Faculty of Medicine, Imperial College London, London, United Kingdom

**Keywords:** Host-pathogen interactions, comparative proteomics, influenza virus polymerase, replication, transcription, antiviral targets

## Abstract

The influenza A virus polymerase, consisting of a heterotrimer of three viral proteins, carries out both transcription and replication of the viral RNA genome. These distinct activities are regulated by viral proteins that vary in abundance during infection, and by various co-opted host cell proteins, which serve as targets for the development of novel antiviral interventions. However, little is known about which host proteins direct transcription and which replication. In this report, we performed a differential interactome screen to identify host proteins co-opted as either transcription-or replication-specific factors. We found that distinct sets of host proteins interact with the influenza polymerase as it carries out the different activities.

We functionally characterised HMGB2 and RUVBL2 as replication-specific cofactors and RPAP2 as a transcription-specific cofactor. Our data demonstrate that comparative proteomics can be used as a targeted approach to uncover virus-host interactions that regulate specific stages of the viral lifecycle.

## Introduction

As an obligate intracellular parasite, influenza A virus (IAV) is heavily reliant on host cell proteins for all steps of its lifecycle, including viral RNA synthesis (1). Understanding which host cell proteins are hijacked and the details of how they support viral activity may improve antiviral strategies with the potential to reduce the public health burden associated with IAV and prevent future IAV pandemics from emerging (2, 3).

The IAV RNA-dependent RNA polymerase catalyses either transcription or bidirectional replication of the viral RNA (vRNA) genome by assuming different complexes that depend on its interaction with various viral or host proteins (1). This happens in the context of viral ribonucleoprotein complexes (vRNPs), in which the vRNA is bound by nucleoprotein (NP) and a polymerase heterotrimer consisting of PB2, PB1, and PA proteins (4). Transcription occurs in *cis*, converting vRNA into viral mRNA, and depends on interaction with the host RNA polymerase (RNAP II) (5, 6). In contrast, replication is a two-step process that relies on the acidic nuclear phosphoprotein 32 (ANP32) family of host proteins (7–11). In the first step, a full-length positive-sense complementary RNA (cRNA) intermediate is synthesised from the vRNA template and packaged into a cRNP (7, 12). In the second step, cRNA is used as a template to produce new vRNA genomes that can also serve as templates for secondary transcription (7). ANP32 is essential for replication as it facilitates assembly of nascent RNA into RNPs (13, 14). It achieves this by stabilising an asymmetric dimer between the replicating polymerase and a newly synthesised polymerase, poised to bind nascent RNA (13). Meanwhile, ANP32 also uses its C-terminal tail to recruit the NP that packages RNA into RNPs (14). Other factors aside from ANP32 have also been found to regulate replication, such as the viral NS2 protein (15, 16). However, it is not known which further host proteins are recruited to the replication platform to promote cRNA and/or vRNA synthesis.

Earlier proteomic and genetic screens have identified host proteins important for the IAV lifecycle (2, 17–27). Most of these studies aimed to generate a global map of host proteins involved in IAV infection using various strategies such as affinity purification (AP) of tagged viral proteins and genome-wide RNA interference (RNAi) or CRISPR screening.

Amongst the many screens, there is little overlap in the host proteins identified and few of the host proteins have been assigned specific roles in the IAV lifecycle (28). Some AP mass spectrometry (AP-MS) screens have focused on the interactome of the IAV polymerase, but such studies often do not distinguish between the host proteins required for transcription versus replication (17–19, 21).

Here, we used cells with ANP32 proteins ablated (ANP32-KO) as a unique tool to uncouple viral transcription and replication, since the latter is blocked in the absence of this host protein. We compared the PB2 interactome in WT and ANP32-KO cells using a live-virus AP-MS strategy and identified host proteins that interacted with either the viral transcriptase or replicase. Using various gene depletion methods and experimental conditions that uncouple cRNA and mRNA synthesis, we showed that the IAV polymerase utilises distinct sets of host proteins for transcription and replication, with HMGB2 and RUVBL2 being examples of replication-specific cofactors and RPAP2 being an example of a transcription-specific cofactor. Together, these data suggest our differential AP-MS strategy to be a more targeted approach to discovering virus-host interactions and understanding their role.

## Results

### The interactome of PB2 is different in WT cells and ANP32-KO cells

The systematic identification of cellular factors that support IAV replication can provide valuable insights into IAV biology, with the potential to inform antiviral strategies. Previous work has shown ANP32 to be an essential cofactor for viral RNA replication, but not transcription (8, 10, 11). Consequently, only steps upstream of RNA replication (virus entry, nuclear import, primary transcription, and translation) are supported in ANP32-KO cells (with ANP32A, ANP32B, and ANP32E ablated) (Figure S1A). To identify host proteins that are co-opted by IAV as replication cofactors, we compared the interactome of PB2 in WT cells, where both viral transcription and replication take place, to the interactome in ANP32-KO cells, where replication does not occur. We reasoned that host proteins co-precipitating from WT, but not ANP32-KO cells, may be polymerase interactors with roles downstream of transcription, some of which may be specific to polymerase replicase activity. Meanwhile, proteins co-precipitating from both cell types may have roles specific to polymerase transcriptase activity.

To study the polymerase interactome in the context of live virus, we generated a recombinant H1N1 A/Puerto Rico/8/1934 (PR8) influenza virus that expresses a FLAG tag on the C-terminus of the PB2 polymerase subunit by reverse genetics (PR8 PB2-FLAG). Virus growth of PR8 PB2-FLAG was similar to that of PR8 PB2-WT (Figure S1B). Retention of the FLAG tag was confirmed by sequencing after virus rescue and one passage (data not shown).

We determined the optimal timepoint post infection for distinguishing replication and transcription host factors based on the accumulation of segment 4 (Haemagglutinin (HA)) vRNA, cRNA, and mRNA, quantified using a tagged RT-qPCR method, in eHAP WT and ANP32-KO cells infected with PR8 PB2-FLAG (Figure 1A-C). eHAP cells were selected as these are amenable to genetic manipulation and are susceptible to IAV infection (9, 29). As expected, no vRNA or cRNA products accumulated above input in the ANP32-KO cells over time (Figure 1A and B). At 2 hours post infection (hpi), mRNA levels had increased 30-fold in both cell types, while cRNA but not yet vRNA had accumulated in the WT cells (Figure 1A-D). We selected the early timepoint of 3hpi for interactome analysis because at this timepoint primary transcription had already taken place in both cell types and cRNA and vRNA synthesis had occurred in the WT cells.

**Figure 1.**
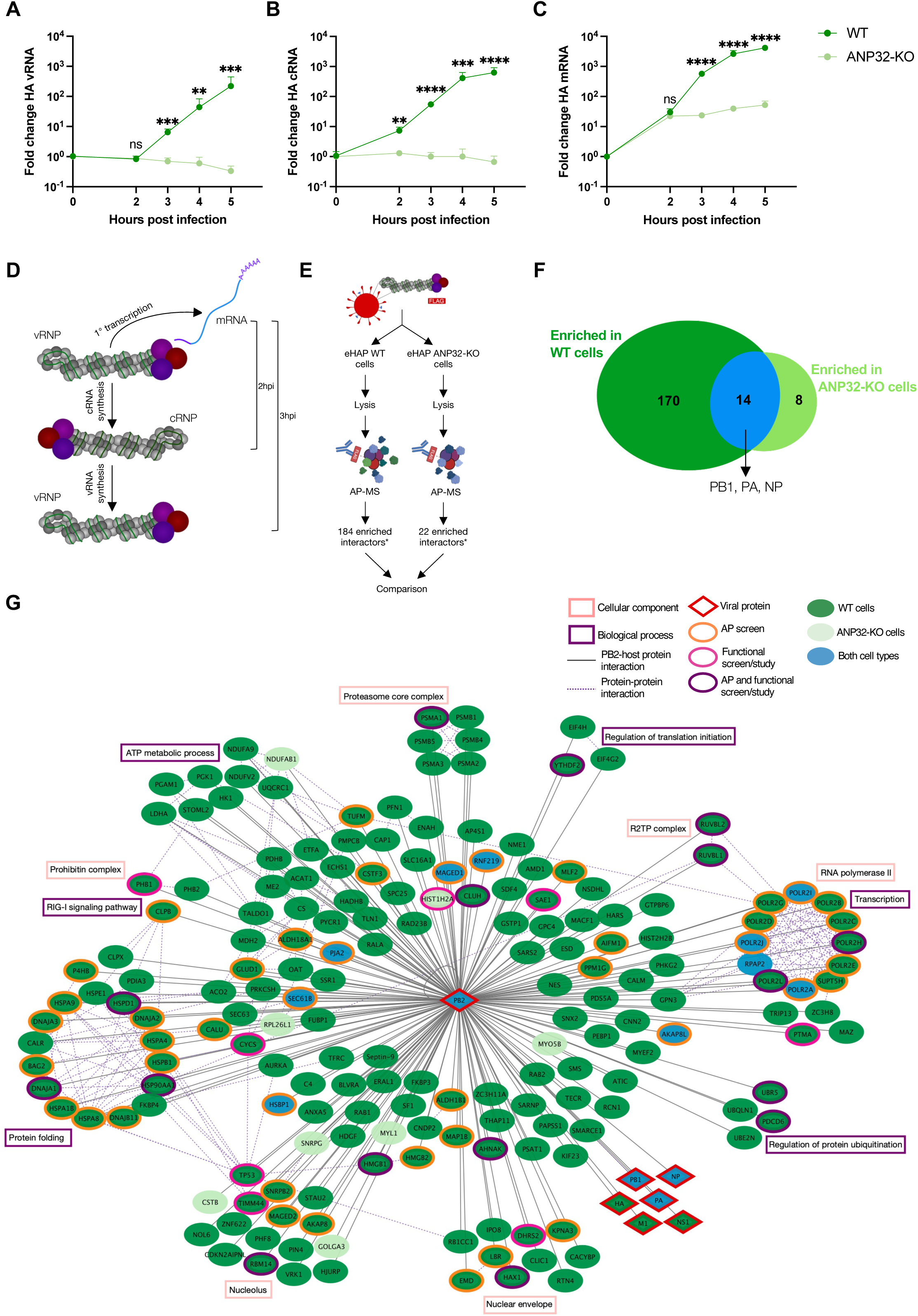
The interactome of PB2 is different in WT cells and ANP32-KO cells. **A-C)** Accumulation of segment 4 (Haemagglutinin (HA)) vRNA (A), cRNA (B), mRNA (C) in eHAP WT and ANP32-KO cells following infection with PR8 PB2-FLAG (MOI=5). The fold change was calculated versus input (0hpi). Data is shown as mean ± SD of n=3 technical repeats, representative of n=3 independent biological repeats. Statistical significance was assessed following log transformation using multiple unpaired *t* tests, corrected for multiple comparisons using the FDR. ns, not significant; **, p<0.01; ***, p<0.001; ****, p<0.0001. **D)** Schematic showing vRNA, cRNA, and mRNA synthesis at 2 and 3hpi in WT and ANP32-KO cells. At 3hpi, primary transcription has occurred in both cell types and vRNA and cRNA synthesis has occurred in WT cells. **E)** Schematic of AP-MS strategy to compare the interactome of PB2 in WT and ANP32-KO cells in the context of infection. * = number of proteins enriched above background (PR8 PB2-WT) in each cell type. **F)** Venn diagram comparing the number of proteins enriched above background in WT cells, ANP32-KO cells, or both WT and ANP32-KO cells. **G)** Interaction network between PB2 (bait, blue) and 170 human proteins in WT cells (green), 8 proteins in ANP32-KO cells (light green), and 13 proteins in both cell types (blue). Viral proteins are diamond-shaped and outlined in red. Human proteins identified in previous selected IAV screens and studies are indicated by outline colour (orange, AP-MS screen; pink, functional screen or study; purple, both AP-MS and functional screen or study). Proteins are clustered based on GO biological process (purple square) or cellular component (coral square) terms. Protein interaction network was generated using Cytoscape v.3.10.2. See also Figure S1 and Table S1 and S2.

To uncover PB2 interaction partners, we infected eHAP WT and ANP32-KO cells with PR8 PB2-FLAG, immunoprecipitated PB2-FLAG, and identified enriched co-precipitating host and viral proteins by MS (Figure 1E). Screens were conducted three independent times and showed good reproducibility with an average Pearson correlation coefficient of 0.86 between each screen repeat. We also ensured comparable lysate amounts of the PB2-FLAG bait in the WT and ANP32-KO cell conditions by using three times more lysate in the latter condition (Figure S1C). Host and viral proteins were classified as interactors based on their enrichment (S_0_=0.2, FDR=0.05) over negative background control (proteins precipitated from WT or ANP32-KO cells infected with WT virus, which lacks a FLAG tag) (Figure S1D and E, Table S1). The negative background controls from either cell type were similar with an average coefficient of variation of 5%.

Using this criteria, 170 host cell proteins were uniquely co-precipitated from WT cells, 8 host proteins from ANP32-KO cells, and 14 proteins were common to both cell types (Figure 1F and S1F). Similar amounts of PB2-FLAG bait were precipitated in both WT and ANP32-KO cells (Figure S1F). Furthermore, the PB1 and PA polymerase subunits and NP were enriched above background in both cell types, confirming that PB2-FLAG was immunoprecipitated in its heterotrimeric and RNP forms (Figure 1F). The higher number of proteins co-precipitating from WT cells was expected, given that all steps following transcription, including replication and vRNP assembly and transport, can occur in WT cells but not in ANP32-KO cells (Figure S1A). In support of this, one of the host proteins uniquely co-precipitating from WT cells was CLUH, which has been shown to promote subnuclear transport of progeny vRNPs (Figure 1G) (30). Also, as expected, less NP co-precipitated from ANP32-KO cells than WT cells, which is consistent with that viral proteins are synthesised from the incoming vRNA template in ANP32-KO cells, but progeny vRNPs do not assemble (Figure S1F).

Our interaction network recapitulated previously identified polymerase interactors and IAV cofactors (Figure 1G, Table S2). For instance, we found PB2 to interact with proteins involved in protein folding, including HSP90 and DNAJA1, which have previously been implicated in the IAV lifecycle (31, 32). Such proteins were uniquely enriched in the WT cells, suggesting that they contribute to the IAV lifecycle downstream of viral transcription. Another interaction specific to the WT cells was with the R2TP complex (RUVBL1 and RUVBL2).

This complex was previously reported to interact with PB1 and RUVBL2 was also reported to interact with vRNPs using NP as bait (17, 20, 22, 33). In addition, many of the PB2 interactors, such as RBM14, are components of the nucleolus. This nuclear site is believed to be important for proper NP assembly and, by extension, RNP formation (34). We also found that PB2 interacted with RNAP II subunits, including the large POLR2A subunit, which is known to be essential for the cap-snatching step of viral transcription (6). This interaction occurred in both WT and ANP32-KO cells, consistent with that viral transcription occurs in both cell types. Our screen also revealed novel interactors such as the FKBP3 and FKBP4 proteins, which were enriched in the WT cells, and RPAP2, which was common to both cell types.

To validate our screen’s ability to identify PB2 interactors, we performed co-immunoprecipitations with two host proteins specifically enriched in the WT cells: HMGB2 and RUVBL2 (Figure S2A-D). WT and ANP32-KO cells were transfected with HMGB2-FLAG or RUVBL2-FLAG prior to WT PR8 infection and FLAG immunoprecipitation. As for the AP-MS screen, we ensured comparable amounts of PB2 in the input from WT and ANP32-KO cells by using three times more ANP32-KO cell lysate than WT cell lysate. Consistent with the interactome data, PB2 co-precipitated with both HMGB2 and RUVBL2 and this occurred to a greater extent in WT cells than in ANP32-KO cells (Figure S2A and C). Considering the small difference in PB2 inputs between the two cell types, we quantified and normalized the PB2 amount in the immunoprecipitate to that in the input (Figure S2B and D). This further corroborated that more PB2 co-precipitated with HMGB2 and RUVBL2 in WT cells than ANP32-KO cells.

### siRNA functional validation reveals host proteins with proviral and antiviral roles in IAV infection

A panel of 13 host proteins uniquely co-precipitating from WT cells, some of which may be important for the polymerase replicase activity, were prioritised for onwards validation based on their host function, nuclear localisation, previous reports of polymerase subunit interaction or known role in the IAV lifecycle (Figure 1G, Table S2). To functionally validate a role for the selected interactors in IAV infection, we assessed the effect of their transient knockdown on virus replication. We made use of a recombinant PR8 virus that expresses a ZsGreen fluorophore on segment 8 (NS) (PR8 NS-ZsGreen) (35). The ZsGreen virus produces three proteins from segment 8: NS1, ZsGreen, and NS2. A549 cells were infected with PR8 NS-ZsGreen after transfection with siRNA pools targeting each of the selected genes, a non-targeting siRNA control (siNT), or two separate siRNA pools targeting both ANP32A and ANP32B (siANP32.1 and siANP32.2) as positive controls for proviral host factors. The impact of each gene knockdown on virus replication was determined by imaging and quantification of ZsGreen at 48hpi (Figure 2A). As expected, knockdown of ANP32A and B significantly reduced virus replication compared to siNT. Knockdown of six of the 13 selected genes (FKBP4, FUBP1, HMGB2, PIN4, RBM14, and RUVBL2) also resulted in a significant reduction in virus replication, suggesting these factors to be proviral. In contrast, knockdown of MAZ significantly increased virus replication, suggesting that this factor has an antiviral role. Gene knockdown at the mRNA level was confirmed by qPCR (Figure S3). Cell viability for all factors remained above 90% following gene knockdown, indicating that the effects on viral replication were a result of gene knockdown and not toxicity (Figure 2B).

**Figure 2.**
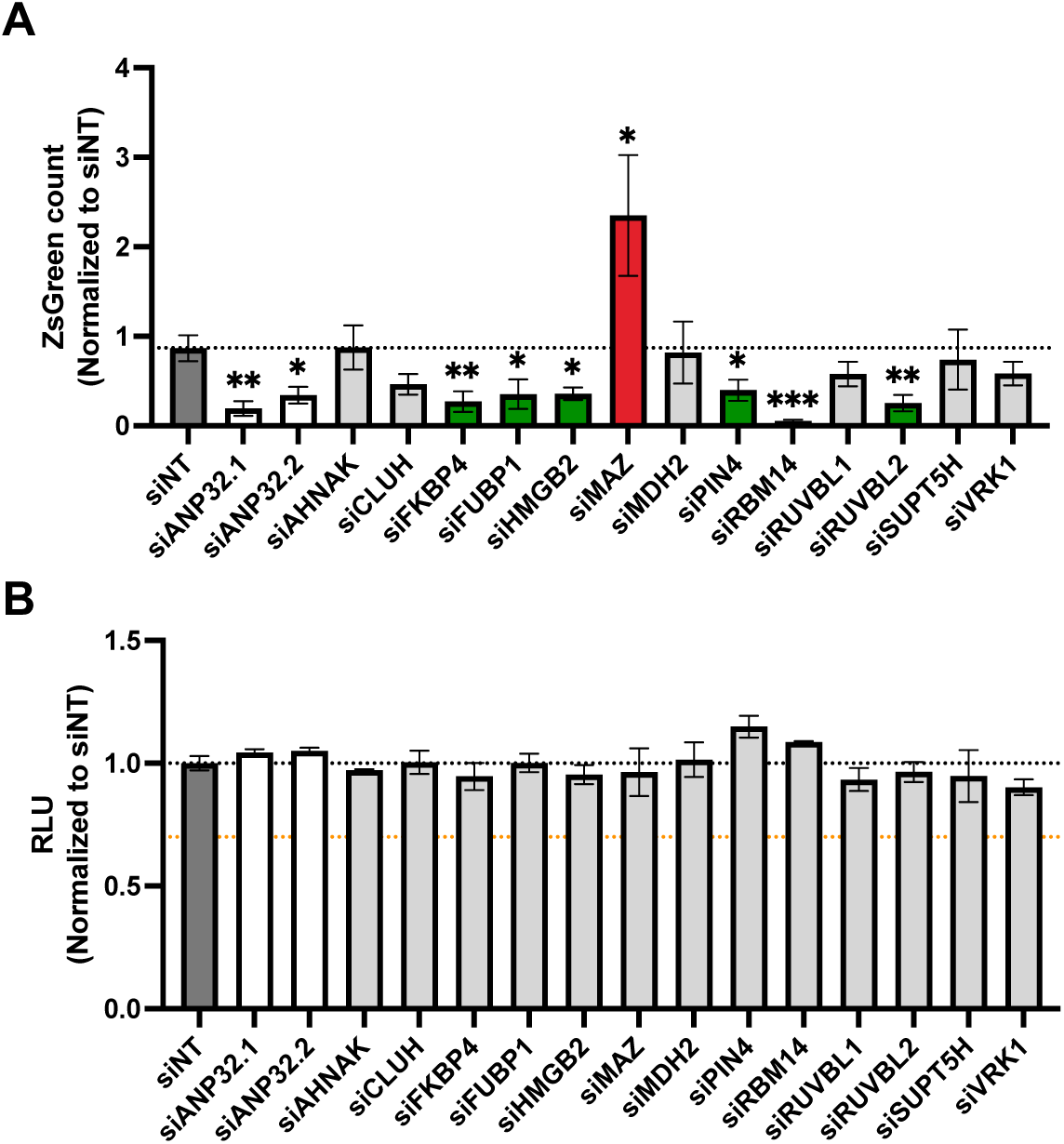
siRNA knockdown reveals host proteins with proviral and antiviral roles in IAV infection. **A)** Virus replication in A549 cells treated with a non-targeting siRNA (siNT), two siRNA pools targeting both ANP32A and ANP32B (siANP32.1 and siANP32.2), or siRNA pools targeting 13 genes enriched in WT cells only. Cells were imaged at 48hpi with PR8 NS-ZsGreen (MOI=0.1). Entire wells were imaged and ZsGreen count was quantified using ImageJ. Proviral factors are shown in green and antiviral factors are shown in red. Data is shown as mean ± SEM of n=3 biological repeats. **B)** Cell viability determined using a CellTiter-Glo assay and shown as luciferase activity (RLU) normalized to siNT. Black dotted line indicates cell viability of siNT-treated cells. Orange dotted line indicates 70% cell viability threshold. Data is shown as mean ± SD of n=3 technical repeats, representative of n=3 biological repeats. For all panels: Statistical significance was assessed compared to siNT using an unpaired *t* test. *, p<0.05; **, p<0.01; ***, p<0.001. See also Figure S3.

Collectively, this revealed seven regulators of IAV infection, six of which were proviral, one of which was antiviral, and two of which (RBM14 and RUVBL2) have previously been identified as host factors involved in IAV infection (22, 33, 36).

### HMGB2, RUVBL2, and RPAP2 promote IAV polymerase activity in human cells

Given that the AP-MS screen was focused on the interactome of PB2 and the main function of PB2 is as part of the heterotrimeric viral polymerase complex, we next sought to confirm a functional role for the interactors in polymerase activity. We decided to further characterise the host factors functionally validated in Figure 2 as well as 18 other host proteins uniquely enriched in the PB2 interactome in WT cells, some of which we hypothesised to be replication-specific host factors (Figure 1G, Table S2). We also characterised two host proteins enriched in both WT and ANP32-KO cells (AKAP8L and RPAP2), which we hypothesised to be transcription-specific host factors (Figure 1G, Table S2). Again, host proteins were selected for onwards validation based on their host function and nuclear localisation as well as previous association with the IAV lifecycle. eHAP WT cells were transfected with siNT, siANP32.1, siANP32.2 or siRNA pools targeting each of the selected genes. Polymerase activity was then measured using a minigenome reporter assay, which requires both replication and transcription of the viral-like reporter RNA to give a luciferase signal (Figure 3A). Efficient gene knockdown, confirmed by qPCR at the mRNA level, did not reduce cell viability to below 80% compared to siNT (Figure S4A and B).

**Figure 3.**
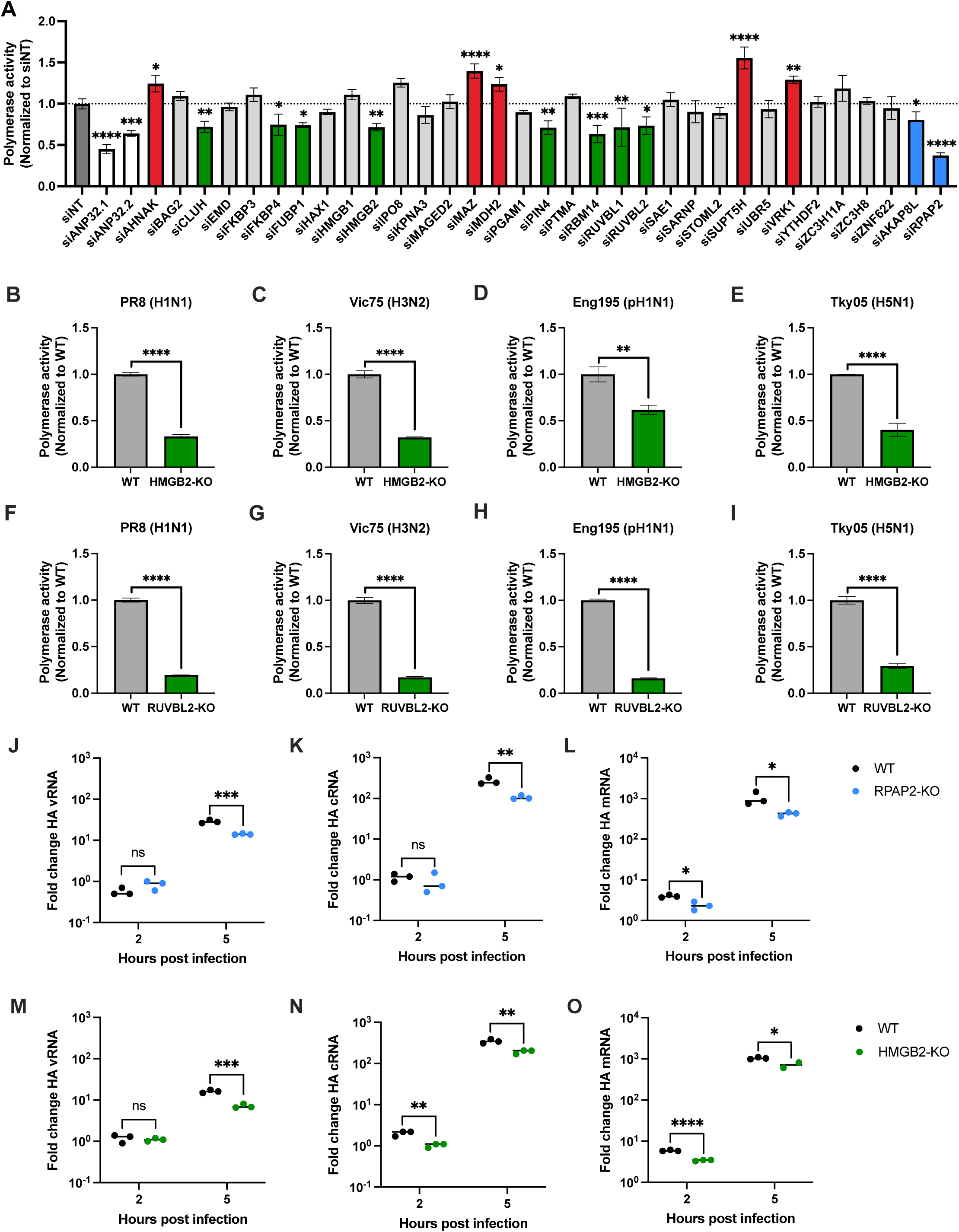
HMGB2, RUVBL2, and RPAP2 promote IAV polymerase activity. **A)** Minigenome reporter assay in eHAP WT cells transfected with a non-targeting siRNA (siNT), two siRNA pools targeting both ANP32A and ANP32B (siANP32.1 and siANP32.2), or siRNA pools targeting 31 genes enriched in WT cells or 2 genes enriched in both WT and ANP32-KO cells. Cells were transfected with plasmids to reconstitute PR8 polymerase activity. Firefly luciferase activity was normalized to co-transfected Renilla luciferase levels. Putative replication-specific proviral factors are shown in green, replication-specific antiviral factors in red, and transcription-specific proviral factors in blue. Statistical significance was assessed compared to siNT using an unpaired *t* test. **B-I)** Minigenome reporter assay in A549 WT and HMGB2-KO cells (B-E) or WT and RUVBL2-KO cells (F-I) transfected with plasmids to reconstitute PR8 (B and F), Vic75 (C and G), Eng195 (D and H), and Tky05 (E and I) polymerase activity. Firefly luciferase activity was normalized to co-transfected Renilla luciferase levels. Statistical significance was assessed compared to WT using an unpaired *t* test. **J-O)** Accumulation of segment 4 (Haemagglutinin (HA)) vRNA (J and M), cRNA (K and N), and mRNA (L and O) in A549 WT and RPAP2-KO cells (J-L) or WT and HMGB2-KO cells (M-O) following infection with PR8 (MOI=3). Data is shown as the fold change over input (0hpi). Statistical significance was assessed at each timepoint following log transformation using multiple unpaired *t* tests, corrected for multiple comparisons using the FDR. For all panels: Data is shown as mean of n=3 technical repeats, representative of n=3 biological repeats. ns, not significant; *, p<0.05; **, p<0.01; ***, p<0.001; ****, p<0.0001. See also Figure S4 and S5.

Knockdown of ANP32A and B resulted in a significant loss of polymerase activity (Figure 3A). Similarly, knockdown of ten of the genes (CLUH, FKBP4, FUBP1, HMGB2, PIN4, RBM14, RUVBL1, RUVBL2, AKAP8L, and RPAP2) significantly reduced polymerase activity compared to siNT, suggesting that these factors have supportive roles in polymerase activity. Of these, six had been found to support viral infection (Figure 2A). Polymerase activity increased following knockdown of five of the genes (AHNAK, MAZ, MDH2, SUPT5H, and VRK1), suggesting that these factors have restrictive roles in polymerase activity (Figure 3A). Of these, MAZ had also shown a restrictive role in the context of live virus (Figure 2A).

As a proof-of-principle that the differential interactome analysis can be used as a tool to distinguish host factors that support either replication or transcription, we decided to further characterise the roles of HMGB2 and RUVBL2 as potential replication-specific cofactors, and RPAP2 as a potential transcription-specific cofactor.

To further investigate the role of HMGB2, RUVBL2, and RPAP2 in polymerase activity, we generated A549 cells lacking expression of HMGB2, RUVBL2, or RPAP2 (HMGB2-KO, RUVBL2-KO, or RPAP2-KO) using CRISPR-Cas9 genome editing. For each gene, we used a pool of three gRNAs targeting an early exon of the gene. A control cell line (WT) was also generated through identical treatment with a non-targeting gRNA. We confirmed that each HMGB2-KO, RUVBL2-KO, and RPAP2-KO clone had an insertion or deletion resulting in a frameshift by Sanger sequencing (Figure S5A-C, data not shown). Western blot analysis revealed that HMGB2 and RUVBL2 protein expression levels were reduced in the HMGB2-KO and RUVBL2-KO clones, respectively, but not completely abrogated (Figure S5D and E). We were unable to detect RPAP2 by western blot.

Activity of reconstituted PR8 polymerase was significantly reduced in the HMGB2-KO and RUVBL2-KO edited cell lines (Figure 3B and F). Polymerase activity was normalized for transfection efficiency using a co-transfected plasmid encoding Renilla luciferase under a RNAP II promoter. As RPAP2 has a host function important for RNAP II transcription activity (37), we were unable to acquire reliable Renilla values from the RPAP2-KO cells and, therefore, minigenome reporter assay data is not reported from these cells. To validate that the reduction in polymerase activity in HMGB2-KO and RUVBL2-KO cells was not due to off-target effects, we reconstituted the HMGB2-KO and RUVBL2-KO cells with exogenously expressed HMGB2 and RUVBL2 and performed minigenome assays to test for complementation (Figure S5F and G). Polymerase activity was only partially restored with HMGB2 complementation (Figure S5F). Rescue of polymerase activity was greatest when HMGB2 expression was similar to HMGB2 levels in WT cells (40ng HMGB2). This suggests that the IAV polymerase is only supported when HMGB2 is expressed at endogenous levels and that too high amounts of HMGB2 may exert an inhibitory effect. Complementation with RUVBL2 fully restored polymerase activity to WT levels in a dose-dependent manner (Figure S5G).

Next, to understand whether the proviral roles of HMGB2 and RUVBL2 were conserved across IAV strains, we measured their impact on activity of the polymerase complex from a seasonal H3N2 strain (A/Victoria/3/75 (Vic75)), a pandemic H1N1 strain (A/England/195/2009 (Eng195)), and a H5N1 strain (A/turkey/Turkey/05/2005 (Tky05)) (Figure 3C-E and G-I). The level of PB2 expression was similar in WT and HMGB2-KO or RUVBL2-KO cells for all strains (Figure S5H and I). Activity of all the viral polymerases tested was significantly reduced in HMGB2-KO and RUVBL2-KO cells, indicating that HMGB2 and RUVBL2 proteins promote polymerase activity of a diverse range of IAV strains (Figure 3C-E and G-I).

As we were unable to acquire minigenome data from the RPAP2-KO cells, we instead investigated the role of RPAP2 in polymerase activity by quantifying the accumulation of segment 4 vRNA, cRNA, and mRNA in WT and RPAP2-KO cells following infection (Figure 3J-L). At 5hpi, a significant reduction in vRNA, cRNA, and mRNA accumulation was observed in RPAP2-KO cells compared to WT cells. This suggests that RPAP2 is important for robust polymerase activity in the context of infection. In similar experiments, we analysed the accumulation of vRNA, cRNA, and mRNA in WT and HMGB2-KO cells following infection (Figure 3M-O). A significant reduction in vRNA, cRNA, and mRNA accumulation was observed in HMGB2-KO cells, which is consistent with our minigenome data and provides further evidence in support of a proviral role for HMGB2 in viral RNA synthesis. Taken together, these data collected in two different cell types (eHAPs and A549s) and using two gene depletion methods (siRNA and CRISPR-Cas9) as well as two methods of measuring polymerase activity (minigenome reporter assay and RT-qPCR) show HMGB2, RUVBL2, and RPAP2 to be proviral host factors that support IAV polymerase activity.

### HMGB2 and RUVBL2 are replication-specific cofactors and RPAP2 is a transcription-specific cofactor

The unique strategy of our AP-MS screen was designed to discover host proteins important for replicase or transcriptase activity. Based on the comparative interactomes obtained in WT and ANP32-KO cells, we hypothesised that HMGB2 and RUVBL2 would be replication-specific cofactors whereas RPAP2 would be a transcription-specific cofactor. To provide evidence that this is the case, we employed assays that allow cRNA and mRNA synthesis to be separated.

First, we quantified the accumulation of vRNA, cRNA, and mRNA in a minigenome setup, following transient knockdown of HMGB2, RUVBL2, and RPAP2 (Figure 4A-C).

**Figure 4.**
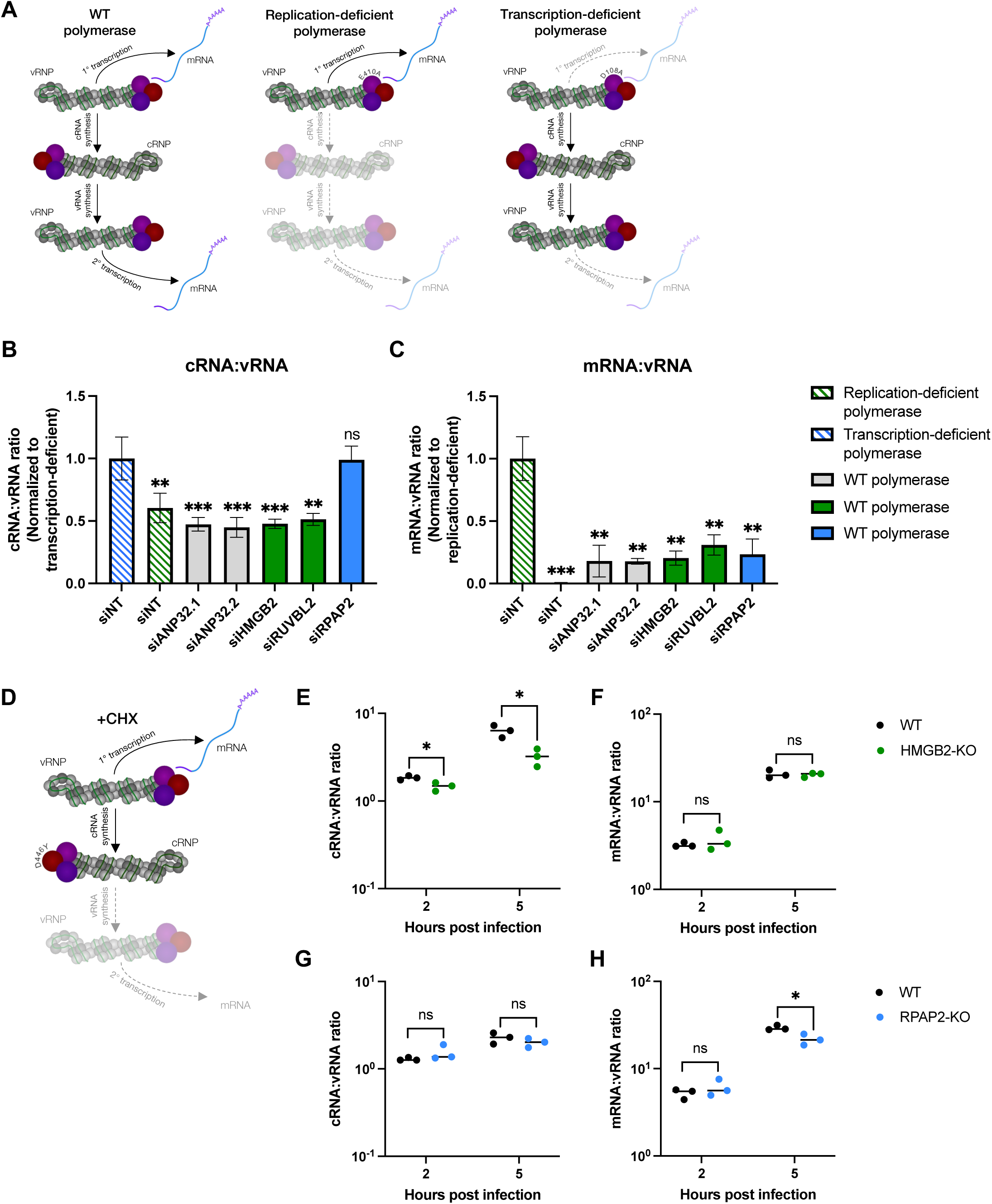
HMGB2 and RUVBL2 are replication-specific cofactors and RPAP2 is a transcription-specific cofactor. **A)** Schematic showing vRNA, cRNA, and mRNA synthesis in a minigenome setup with a WT, replication-deficient (PA E410A), or transcription-deficient (PA D108A) polymerase. **B and C)** cRNA:vRNA ratio (B) and mRNA:vRNA ratio (C) produced by a PR8 WT vRNP following transient knockdown of both ANP32A and ANP32B (siANP32.1 and siANP32.2), HMGB2, RUVBL2, or RPAP2. Activity of a replication-deficient vRNP (green striped) and transcription-deficient vRNP (blue striped) in cells transfected with a non-targeting siRNA (siNT) is also shown. cRNA:vRNA ratios were normalized to the ratio produced by the transcription-deficient vRNP in siNT-treated cells (B). mRNA:vRNA ratios were normalized to the ratio produced by the replication-deficient vRNP in siNT-treated cells (C). vRNA template provided was a pPolI-Firefly luciferase plasmid. Statistical significance was assessed compared to replication-deficient/transcription-deficient polymerases using an unpaired *t* test. **D)** Schematic showing vRNA, cRNA, and mRNA synthesis under conditions of a cRNP stabilization assay with cycloheximide (CHX) treatment. **E-H)** cRNP stabilization assay with CHX in A549 WT and HMGB2-KO cells (E and F) or WT and RPAP2-KO cells (G and H). Data is shown as fold change in segment 4 (Haemagglutinin (HA)) cRNA:vRNA ratio (E and G) and mRNA:vRNA ratio (F and H) over input (0hpi). Statistical significance was determined at each timepoint following log transformation using multiple unpaired *t* tests, corrected for multiple comparisons using the FDR. For (B), (C), and (E-H): Data is shown as mean of n=3 technical repeats, representative of n=3 biological repeats. ns, not significant; *, p<0.05; **, p<0.01; ***, p<0.001. See also Figure S6.

Unlike in infection where vRNA, cRNA, and mRNA synthesis are interdependent, in a minigenome setup, vRNA and cRNA synthesis (replication) occur independently of mRNA synthesis (transcription). We employed replication-deficient (PA E410A) and transcription-deficient (PA D108A) mutant polymerases as controls (Figure 4A) (38). The replication-deficient polymerase generated less vRNA and cRNA than WT polymerase, but near WT levels of mRNA (Figure 4A and S6A-C). Conversely, the transcription-deficient polymerase produced equivalent vRNA and cRNA levels as WT polymerase but lower mRNA levels (Figure 4A and S6A-C). We, therefore, compared the cRNA:vRNA ratio following knockdown of HMGB2, RUVBL2, and RPAP2 to that of the transcription-deficient polymerase and the mRNA:vRNA ratio to that of the replication-deficient polymerase (Figure 4B and C).

Knockdown of ANP32A and B decreased the cRNA:vRNA ratio (Figure 4B). As expected, given that vRNA also acts as a template for mRNA synthesis, the mRNA:vRNA ratio also decreased following knockdown of ANP32A and B (Figure 4C). Knockdown of HMGB2 and RUVBL2 similarly resulted in a decrease in the cRNA:vRNA and mRNA:vRNA ratios (Figure 4B and C). In contrast, knockdown of RPAP2 had no effect on the cRNA:vRNA ratio, but significantly reduced the mRNA:vRNA ratio. Overall, this shows that HMGB2 and RUVBL2 have supportive roles in RNA replication, while RPAP2 has a supportive role in RNA transcription, as we predicted based on the AP-MS differential interactome screen.

To confirm using a different approach that HMGB2 acts as a replication-specific cofactor, whereas support from RPAP2 is transcription-specific, we made use of a previously established cRNP stabilization assay (11, 39) (Figure 4D). A catalytically dead PB1 mutant (PB1 D446Y) was pre-expressed alongside PA, PB2, and NP in cells that were then infected with influenza virus to supply a vRNP template. Pre-expression of this combination of inactive polymerase with NP allows stabilization of cRNA products but not onwards replication. The cells were then treated with cycloheximide (CHX), which blocks translation of any new proteins encoded from the incoming virus, such that only the pioneering round of cRNA synthesis and primary mRNA synthesis can occur.

A significant decrease in cRNA accumulation was observed in HMGB2-KO cells compared to WT cells at 2 and 5hpi (Figure 4E). There was no difference in mRNA accumulation between the two cell types (Figure 4F). This demonstrates that HMGB2 has a direct role in cRNA synthesis, confirming it as a replication-specific cofactor. In contrast, at 5hpi, less mRNA accumulated in RPAP2-KO cells than WT cells, while no difference in cRNA accumulation was observed between the two cell types (Figure 4G and H). This corroborates that RPAP2 has a direct role in mRNA synthesis. Taken together, these data provide further evidence that HMGB2 and RPAP2 are important host factors supporting IAV polymerase replicase or transcriptase activity, respectively, and confirm that the differential AP-MS screen can be used as a tool to identify replication-and transcription-specific host factors.

### Chemical inhibition of HMGB1/2 suppresses virus growth and viral RNA replication

Inflachromene (ICM) is a small molecule inhibitor of HMGB1 and HMGB2, which share 80% amino acid homology (40). ICM binds and prevents phosphorylation and acetylation of the HMGB proteins. Given the direct role of HMGB2 in RNA replication, we sought to investigate the effect of ICM on virus replication and viral RNA synthesis.

ICM concentrations up to 20uM had no effect on the viability of eHAP WT cells compared to the DMSO-treated control (Figure 5A). To assess the effect of ICM on viral replication, eHAP WT cells were infected with WT PR8 and treated with ICM or DMSO (Figure 5B). ICM significantly reduced virus growth compared to the DMSO control. We then assessed the effect of ICM on polymerase activity of two different IAV strains; PR8 and Eng195 (Figure 5C and D). Polymerase activity of both strains decreased in a dose-dependent manner.

**Figure 5.**
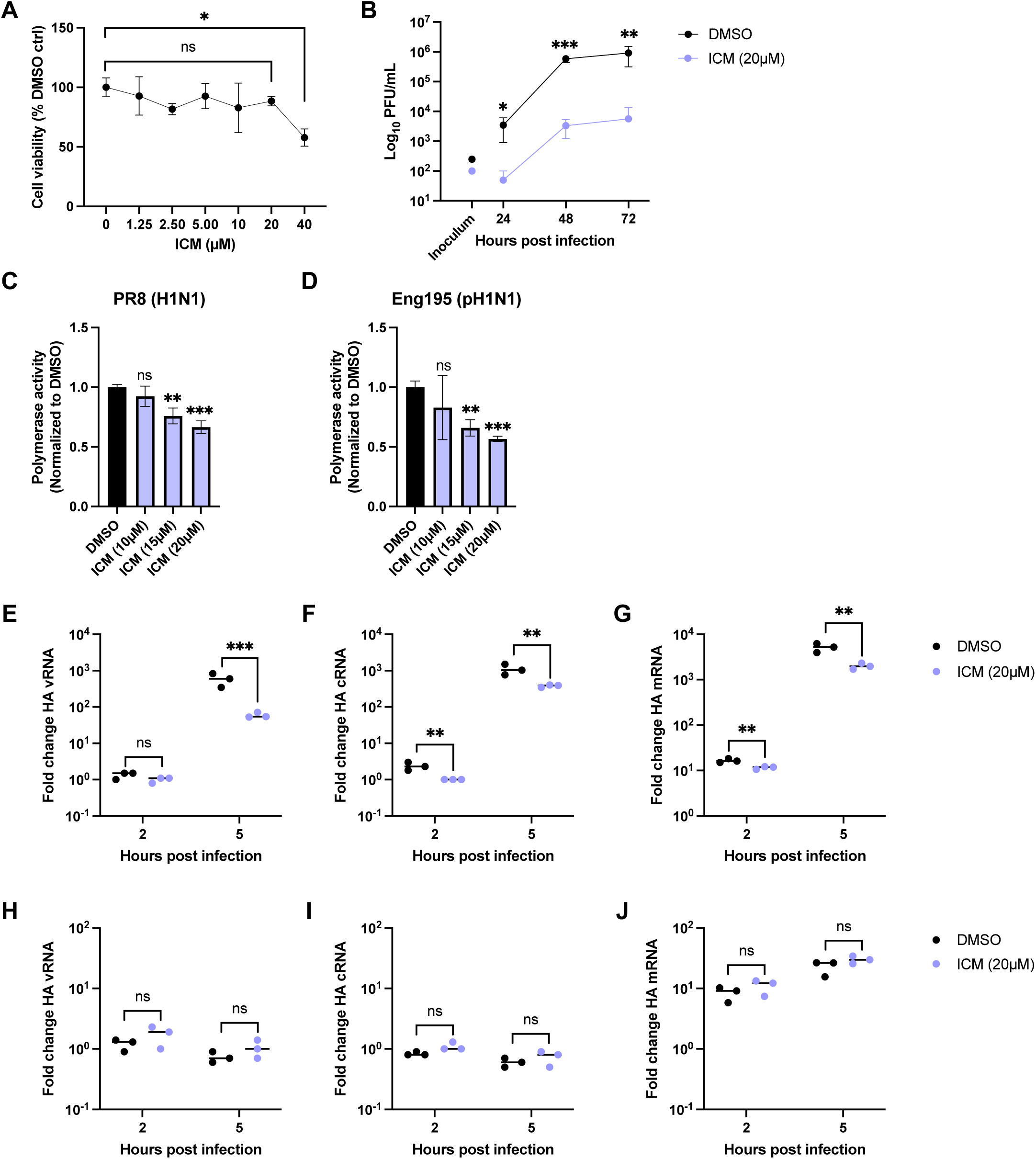
Inflachromene (ICM) suppresses virus growth and viral RNA replication. **A)** Cell viability of eHAP WT cells following ICM treatment (0-40µM) determined using an MTS assay. Cell viability is shown as percentage of DMSO control. Statistical significance was assessed at each concentration using an unpaired *t* test. **B)** Virus replication in eHAP WT cells infected with PR8 (MOI=0.001) and treated with ICM (20µM) or DMSO. Supernatants were harvested at indicated timepoints post infection, and titres were determined by plaque assay on MDCK cells. Statistical significance was assessed at each timepoint using an unpaired *t* test. **C and D)** Minigenome reporter assay in eHAP WT cells treated with ICM (10, 15, or 20µM) or DMSO and transfected with plasmids to reconstitute PR8 (C) or Eng195 (D) polymerase activity. Firefly luciferase activity was normalized to co-transfected Renilla luciferase levels. Statistical significance was assessed compared to DMSO-treated control using an unpaired *t* test. **E-J)** Accumulation of segment 4 (Haemagglutinin (HA)) vRNA (E and H), cRNA (F and I), and mRNA (G and J) in eHAP WT cells (E-G) or ANP32-KO cells (H-J) following infection with PR8 (MOI=3) and treatment with ICM (20µm) or DMSO. Data is shown as the fold change over input (0hpi). Statistical significance was assessed at each timepoint following log transformation using multiple unpaired *t* tests, corrected for multiple comparisons using the FDR. For all panels: Data is shown as mean of n=3 technical repeats, representative of n=3 biological repeats. ns, not significant; *, p<0.05; **, p<0.01; ***, p<0.001.

To further validate that ICM phenocopies the knockout of HMGB2, we compared the accumulation of segment 4 vRNA, cRNA, and mRNA in eHAP WT and ANP32-KO cells infected with PR8 and treated with ICM or DMSO (Figure 5E-J). In eHAP WT cells, ICM reduced the accumulation of all three viral RNA species over time, similar to the reduced viral RNA synthesis observed in HMGB2-KO cells (Figure 5E-G and 3M-O). To understand whether ICM directly inhibits vRNA and cRNA synthesis or indirectly via inhibition of mRNA synthesis, we tested the effect of ICM on the accumulation of viral RNA species in ANP32-KO cells, where only viral transcription can take place (Figure 5H-J). As expected, vRNA and cRNA levels did not increase above input in the ICM-and DMSO-treated ANP32-KO cells, confirming that only mRNA synthesis is supported in these cells (Figure 5H and I). mRNA levels remained the same between ICM-and DMSO-treated ANP32-KO cells, implying that ICM has no effect on transcription and specifically inhibits the cRNA and/or vRNA synthesis steps of replication (Figure 5J). Overall, these data corroborate a direct role for HMGB2 in RNA replication and suggest ICM to be a novel antiviral compound that warrants further investigation.

## Discussion

Many host cell proteins that impact influenza virus have been identified in previous interactome screens, but only a fraction of these have been functionally validated and assigned specific roles in the IAV lifecycle. Here, we build upon previous interactome studies that have shed light on the host proteins co-opted by the IAV polymerase (17–19, 21). We performed a targeted AP-MS screen, subtracting the polymerase interactome when it is in a transcription state (ANP32-KO cells) from its interactome under standard cellular conditions (WT cells). This allowed us to separate the interactome into two categories: 1) host proteins recruited upstream of and for primary transcription and 2) host proteins recruited downstream of and for RNA replication.

We provide functional evidence that our differential interactome analysis can be used to separate the IAV transcriptase and replicase host cell machinery already at the AP-MS screening stage. By further characterising host proteins enriched in both cell types (RPAP2) and in WT cells only (HMGB2 and RUVBL2), we show that all three host factors promote polymerase activity, with RPAP2 doing so by supporting mRNA synthesis and RUVBL2 and HMGB2 doing so by supporting cRNA synthesis.

The protein that we propose supports transcription only is RPAP2. This protein has no previous links to IAV, but the proviral role for RPAP2 in viral transcription shown here fits well with its host function, where it directly binds to RNAP II and supports its nuclear import (37). Interestingly, RPAP2 also plays an additional, more specific role in host transcription, binding the RNAP II CTD phosphorylated at Ser7 and promoting transcription of small nuclear RNA (snRNA) genes (41, 42). snRNA genes are the preferred target for viral cap-snatching, a process in which the IAV polymerase acquires host capped RNA fragments that act as primers for transcription initiation (43). We, therefore, speculate that RPAP2 plays a role in the cap-snatching step of IAV transcription, perhaps bringing the IAV polymerase into proximity with host RNAP IIs transcribing snRNA genes.

We propose that RUVBL2 plays a supportive role in RNA replication. RUVBL2 has previously been identified as a polymerase and NP interactor in various screens, but its role in IAV infection remains unclear (17, 20, 22, 33). A truncated version of quail RUVBL2 was reported to promote IAV polymerase activity, while full-length human RUVBL2 was shown to be a suppressor of polymerase activity when overexpressed (33). More recent work has shown human RUVBL2 to be a proviral factor in IAV infection (22). Our data agrees with this latter study, and we further show RUVBL2 acts at the RNA replication stage of the IAV lifecycle. Considering the essential role of NP in cRNA and vRNA synthesis but not mRNA synthesis, a role for RUVBL2 specifically in RNA replication is consistent with previous reports of a direct interaction between RUVBL2 and free NP (33).

We found that knockdown of RUVBL2 not only reduced cRNA but also mRNA levels. This may be an indirect effect due to the dependency of transcription on replication to amplify the vRNA template. However, we cannot exclude the possibility that RUVBL2 also plays an additional role in viral transcription. Interestingly, in the host cell, RUVBL2 also binds the RNAP II CTD, promoting RNAP II clustering and transcription initiation (44). This could implicate RUVBL2 in supporting viral transcription as well, given the role of host RNAP II in IAV transcription (6). However, whether the IAV polymerase remains associated with or dissociates from host RNAP II during RNA replication is not resolved (45, 46). In the future, it will be interesting to confirm whether RUVBL2 also plays a role in transcription, in addition to its role in RNA replication shown here, and whether its association with host RNAP II plays a role. RUVBL2 has also been shown to interact with the viral NS1 protein and contribute to the regulation of apoptosis in infected cells (47). It is possible that RUVBL2 plays two separate roles in IAV infection, where our study has focused on its role in IAV polymerase activity.

We propose HMGB2 to be a cofactor specific to RNA replication. Despite being reported as an NP interactor in human cells, HMGB2 was previously found to have no effect on virus replication in HMGB2-KO mouse embryonic fibroblasts (48). In further contrast to our data, the authors found HMGB1 to have a supportive role in IAV infection in these cells (48). Considering that our data was acquired from human cells, the contradiction between the findings could be because of differences in the roles of HMGB1 and HMGB2 in mice and humans. Interestingly, HMGB2 exists in the same SET complex as ANP32 in the host cell nucleus (49). Considering the ability of HMGB2 to interact with NP, it is possible that HMGB2 works in conjunction with ANP32 to recruit NP to nascent RNAs and aid assembly of RNPs (14, 48).

We demonstrated the application of HMGB2 as an antiviral drug target using ICM, an inhibitor of HMGB1 and HMGB2 that has previously been studied in the context of neuroinflammatory diseases (40). Our data suggests that ICM is a promising antiviral against IAV, suppressing virus growth by blocking viral RNA replication. In the future, it will be interesting to also test the *in vivo* efficacy of ICM in an animal model of IAV infection. When doing so it will be important to consider the cross-species differences in HMGB1 and HMGB2, given that mouse and human HMGB1/HMGB2 appear to play opposite roles in IAV infection (HMGB1 is proviral in mice, HMGB2 is proviral in humans) (48).

Current anti-IAV drugs target viral proteins, but their long-term effectiveness is limited as resistance to such drugs readily emerges (50). Targeting host factors instead may make it more difficult for the virus to evolve resistance. Understanding the exact stage of the viral lifecycle that host factors targeted by antivirals are involved in is important. This is because it allows multiple drugs that block different stages of the viral lifecycle to be used together, further reducing the risk that antiviral resistance arises. For instance, ICM, which we found to block RNA replication, could be used together with baloxavir, which blocks viral transcription (51). Importantly, ICM also has anti-inflammatory properties (40). It might, therefore, be a particularly effective antiviral in the event of a pandemic where infections often result in overexuberant inflammation and this can lead to severe disease (52).

Further studies are needed to confirm the molecular details behind the roles of RPAP2, RUVBL2, and HMGB2 proposed here. Although we have shown HMGB2 and RUVBL2 to promote RNA replication, it remains unclear whether they promote both cRNA and vRNA synthesis and whether they work independently or together with ANP32.

Considering that HMGB2 and RUVBL2 likely act on RNA replication via their interactions with NP, it is conceivable that they play roles in RNP assembly or NP recruitment and, therefore, support both steps of replication. It is likely that such NP interactors were identified in our AP-MS screen because the screen was performed in the context of active virus. This means that we not only captured direct PB2 interactors but also interactors of PB2-containing complexes, such as polymerase heterotrimers and RNPs. Some of the further host proteins identified in our screen in each category may also eventually prove to be effective antiviral targets.

To conclude, we show that the IAV polymerase interacts with distinct sets of host proteins during RNA transcription and replication. The data presented in this study is a proof-of-principle that comparative proteomics of viral proteins locked in specific functional states can be used to identify host factors more precisely. Fundamentally, the AP-MS strategy could be applicable to any virus to directly categorise virus-host interactions and streamline functional validation of hits and identification of antiviral targets.

## Resource availability

Further information and requests for resources and reagents should be directed to and will be fulfilled by the lead contact, Wendy Barclay.

## Data availability

The mass spectrometry proteomics data have been deposited to the ProteomeXchange Consortium via the PRIDE (53) partner repository with the dataset identifier PXD064673.

## Acknowledgements

We thank Mark Skehel and Fairouz Ibrahim at the Proteomics Science Technology Platform at the Francis Crick Institute for analysis of samples by mass spectrometry. We thank Edward Hutchinson for the kind gift of the PR8 NS-ZsGreen plasmid. We thank Isabel Correa-Otero at the FACS facility at Imperial College London for support with single cell sorting. We also thank Goedele Maertens for helpful discussions throughout. This study was supported by Wellcome Trust grant 205100/Z/16/Z. W.S.B. is supported in part by the NIHR Biomedical Research Centre (BRC) (NIHR203323) of Imperial College NHS Trust. In addition, A.B.R. was supported by a Medical Research Council (MRC) studentship.

## Author contributions

Conceptualization, W.S.B. and A.B.R.; Methodology, W.S.B., L.M.S., C.M.S., A.B.R.; Investigation, A.B.R., O.C.S., K.S., N.L., M.D.B.; Resources, L.M.S.; Writing-original draft, W.S.B. and A.B.R.; Writing-review & editing, L.M.S., C.M.S., M.D.B.; Supervision, W.S.B., O.C.S., C.M.S.; Funding acquisition, W.S.B.

## Declaration of interests

The authors declare no competing interests.

## Supplemental information

**Document S1.** Figure S1-S6.

**Table S1.** Raw and processed mass spectrometry data. Related to Figure 1.

**Table S2.** Prioritisation of host proteins for onwards validation. Related to Figure 1.

**Table S3.** Comparison of host proteins identified in this screen to selected previous screens. Related to STAR methods.

**Table S4.** Oligonucleotide sequences. Related to STAR methods.

## STAR methods

### Experimental model and subject details

#### Cells and cell culture

Human engineered haploid (eHAP; Horizon Discovery) cells and eHAP cells with ANP32A, ANP32B, and ANP32E knocked out by CRISPR-Cas9 (29) were maintained in Iscove modified Dulbecco medium (IMDM; Gibco, Life Technologies) supplemented with 10% fetal bovine serum (FBS; Labtech), 1% non-essential amino acids (NEAA; Gibco, Life Technologies), and 1% penicillin-streptomycin (pen-strep; Gibco, Life Technologies).

Adenocarcinomic human alveolar basal epithelial (A549; ATCC), A549-Dox-Cas9 (54), human embryonic kidney 293T (HEK293T; ATCC), and Madin-Darby canine kidney (MDCK; ATCC) cells were maintained in Dulbecco modified Eagle medium (DMEM; Gibco, Life Technologies) supplemented with 10% FBS (Gibco, Life Technologies), 1% NEAA, and 1% pen-strep. When siRNA transfected, eHAP and A549 cells were maintained in IMDM or DMEM, respectively, supplemented with 20% FBS and 1% NEAA. All cells were cultured at 37°C and 5% CO_2_.

#### Viruses and infections

Influenza PR8-WT, PR8 PB2-FLAG, and PR8 NS-ZsGreen viruses were generated by reverse genetics using the pHW2000 reverse genetics system (55). The Flag epitope was introduced into the coding region of PB2 in the pHW2000 plasmid. It was inserted at the C-terminus followed by a duplication of the packaging signal of 173 nucleotides, as described previously (20). The NS-ZsGreen plasmid was a kind gift from Edward Hutchinson (35).

PR8-WT, PR8 PB2-FLAG, and PR8 NS-ZsGreen viruses were propagated in 10-day old chicken eggs and infectious titres were determined by plaque assay on MDCK cells.

Sequencing of viral stocks was used to confirm the presence of the Flag and ZsGreen tags in the rescued viruses and after one passage.

For infections, virus was diluted in serum-free (SF) media to the correct multiplicity of infection (MOI, as indicated in figure legends). For multi-cycle infections, cells were infected with viral inoculum and incubated at 37°C for 1hr. Inoculum was replaced with prewarmed serum-free (SF) media supplemented with 1µg/mL TPCK trypsin (Worthington Biochemical) and infected cells were incubated at 37°C. At the appropriate timepoint, supernatants were harvested for downstream processing or cells were imaged, as described below. For synchronised single-cycle infections, cells were incubated at 4°C for 15min prior to addition of viral inoculum. After addition of viral inoculum, cells were incubated at 4°C for 45min.

Inoculum was replaced with prewarmed full media and infected cells were incubated at 37°C. At the appropriate timepoint post infection, cells were lysed in buffer RLT (Qiagen) supplemented with 10% β-mercaptoethanol (BME; Sigma) or harvested for downstream processing, as described below.

### Method details

#### Flag-tag affinity purifications

eHAP WT cells were cultured in 1×15cm and 1×10cm dishes and eHAP ANP32-KO cells were cultured in 4×15cm dishes. For MS analysis, cells were infected with PR8 PB2-FLAG or PR8-WT at MOI=5, as described above. For immunoblot analysis, cells were transfected with pCAGGS plasmids encoding HMGB2-FLAG (10µg) or RUVBL2-FLAG (5µg) using Lipofectamine 3000 (Invitrogen), 20hrs prior to infection with PR8-WT (MOI=5). Cells were harvested at 3hpi, washed x3 in ice-cold phosphate-buffered saline (PBS; Gibco, Life Technologies), and lysed in lysis buffer (50mM Tris-Cl (pH8.0), 200mM NaCl, 1mM EDTA, 0.2% NP40) supplemented with EDTA-free protease inhibitor cocktail tablet (Roche) on a rotating wheel at 4°C for 30min. Lysates were centrifuged at 13000xg for 10min.

Supernatants were incubated with Anti-FLAG magnetic beads (ThermoFisher Scientific), equilibrated in tris-buffered saline (TBS), on a rotating wheel at 4°C overnight. For MS analysis, beads were washed x4 in lysis buffer, x4 in TBS, and x7 in 50mM ammonium bicarbonate and then analysed by MS, described below. For immunoblot analysis, beads were washed x4 in lysis buffer and x4 in TBS prior to elution by competition with 3xFLAG peptide (ThermoFisher Scientific) for 30min at room temperature (RT). Supernatants were mixed with 4xLaemmli sample buffer (Bio-Rad) with 10% BME and then analysed by immunoblotting, described below.

#### Mass spectrometry

Bead-bound proteins were prepared for mass spectrometric analysis by in solution enzymatic digestion. Briefly, bead-bound proteins in 50mM ammonium bicarbonate were reduced in 10mM DTT and then alkylated with 55mM iodoacetamide. 0.5µg of Trypsin (ThermoFisher Scientific) was added, and the proteins digested for 1hr at 37°C in a thermomixer (Eppendorf), shaking at 800rpm. Following this initial digestion, a further 0.5µg of Trypsin (ThermoFisher Scientific) was added, and digestion continued overnight at 37°C. 1µl of formic acid was added and the beads centrifuged for 60s at 2000rcf, before placing them in a magnetic rack. The supernatant was then removed into a fresh, labelled tube. The resulting peptides were analysed by nano-scale capillary LC-MS/MS using an Ultimate U3000 HPLC (ThermoScientific Dionex) to deliver a flow of approximately 250nL/min. A C18 Acclaim PepMap100 5µm, 75µm x 20mm nanoViper (ThermoScientific Dionex), trapped the peptides prior to separation on a C18 Acclaim PepMap RSLC 3µm, 75µm x 500mm nanoViper (ThermoScientific Dionex). Peptides were eluted with a 90min gradient of acetonitrile (2%v/v to 80%v/v). The analytical column outlet was directly interfaced via a nano-flow electrospray ionisation source, with a hybrid dual pressure linear ion trap mass spectrometer (Orbitrap Lumos; ThermoScientific). Data dependent analysis was carried out, using a resolution of 120,000 for the full MS spectrum, followed by ten MS/MS spectra in the linear ion trap. MS spectra were collected over a m/z range of 300-1500. MS/MS scans were collected with the standard AGC target, dynamic maximum injection time mode, isolation window at 1.2 m/z and 32% normalised HCD collision energy. All raw files were processed with MaxQuant 2.4.9.0 (56) using standard settings and searched against a UniProt Human Reviewed KB/Influenza concatenated database with the Andromeda search engine (57) integrated into the MaxQuant software suite. Enzyme search specificity was Trypsin/P for both endoproteinases. Up to two missed cleavages for each peptide were allowed.

Carbamidomethylation of cysteines was set as fixed modification with oxidized methionine and protein N-acetylation considered as variable modifications. The search was performed with an initial mass tolerance of 6ppm for the precursor ion and 0.5Da for MS/MS spectra. The false discovery rate was fixed at 1% at the peptide and protein level.

#### Mass spectrometry analysis

Statistical analysis was carried out using the Perseus module (v2.0.11) of MaxQuant (58). Prior to statistical analysis, peptides mapped to known contaminants, reverse hits, and protein groups only identified by site were removed. Only protein groups identified with at least two peptides, one of which was unique, and two quantitation events were considered for data analysis. Missing values were imputed using normal distribution, with standard deviation (SD) of 30% and the mean offset by-1.8 SD. To eliminate non-specific background binders, the significance of the protein enrichment in immunoprecipitations with PB2-FLAG as bait versus control experiments (immunoprecipitations from cells infected with WT virus (no FLAG tag)) for each cell type was determined using a *t* test (two-sided, equal variance, s_0_=0.2) and corrected for multiple hypotheses using FDR (FDR=0.05). A similar method of determining protein enrichment was described in (21). Proteins identified in the FLAG CRAPome (59) were manually removed. The remaining proteins enriched in WT cells were compared to the remaining proteins enriched in ANP32-KO cells.

#### Interaction network and gene ontology

The protein interaction network was visualised in Cytoscape (v.3.10.2) (60). Host protein-protein interactions were predicted using STRING with high confidence (score >0.9) (61). The PB2 interactors were analysed for pathway and gene set enrichment using gene ontology (GO) biological process and GO cellular component terms. The GO terms were investigated and manually curated based on published literature. Genes that were part of enriched biological process or cellular component terms were manually clustered together and labelled. The cell type (WT cells, ANP32-KO cells, both cell types) that the virus-host protein interaction occurred in was reported by the node fill colour. The PB2 interactors were compared to host proteins identified in previous AP-MS screens. A range of AP-MS screens using either PB2, PB1, or PA as bait in the context of transfection (2, 22) or infection (19–21) were selected for comparison. To investigate the functional relevance of the PB2 interactors, comparisons were also made to functional screens (23–26) and studies (31–33, 36, 48, 62, 63) (Table S2 and S3). The shared interactors were reported based on node outline colour.

#### siRNA transfection

A549 or eHAP WT cells were transfected with siRNA pools (siGENOME SMARTpool siRNA, Dharmacon) targeting human genes of interest (as indicated in Figure 2, 3, and 4) or a non-targeting siRNA control. siRNA pools targeting both ANP32A and ANP32B were generated by combining individual siRNAs (siGENOME, Dharmacon) targeting ANP32A and ANP32B. Reverse transfections were performed using Lipofectamine RNAiMAX (Invitrogen) according to the manufacturer’s instructions. 48hrs post transfection, transfected cells were used in infection assays (immunofluorescence readout) and minigenome assays (luciferase and RT-qPCR readouts). Gene knockdown at the mRNA level was determined by RT-qPCR using primers specific to each gene and a QuantiNova SYBR green kit (Qiagen) on a Viia 7 real-time PCR system. β-actin was used as an internal control. Primers used are shown in Table S4. Cell viability was assessed using a CellTiter-Glo luminescent viability assay (Promega), according to the manufacturer’s instructions.

#### CRISPR-Cas9 genome editing

Commercial multi-guide sgRNAs targeting HMGB2, RUVBL2, and RPAP2 (Synthego) and non-targeting sgRNA (Synthego) were used. To induce Cas9 expression, A549-dox-Cas9 cells were treated with doxycycline (1µg/mL; Sigma) for 48hrs before transfection with gRNAs using Lipofectamine RNAiMAX, according to the manufacturer’s instructions. 48hrs post transfection, cells were single-cell sorted into 96-well plates containing growth medium using a fluorescence-activated cell sorter (FACS) Aria III (BD Biosciences) with a 100um nozzle. Clonal populations were grown over 14-21 days.

Genomic DNA was extracted using a genomic DNA purification kit (ThermoFisher Scientific). Genetic loci with indels were amplified by PCR and Sanger sequenced. Primers used are shown in Table S4. Sanger sequencing data was analysed using Inference of CRISPR Edits (ICE; Synthego), a software that analyses CRISPR edits and assesses editing efficiency.

Gene knockout at the protein level was confirmed by immunoblotting, described below.

#### Immunofluorescence

Cells were transfected with siRNAs and infected with PR8 NS-ZsGreen, as described above. At 48hpi, cells were imaged on an M5000 EVOS microscope. Entire wells were imaged using the GFP (470/525 nm) light cube, 4x objective, and the automated focus function. ZsGreen count was quantified using ImageJ (64). Briefly, images were converted to greyscale, the threshold was set, and the virus particle count was quantified using the “analyse particle” tool. The parameters were set to the size and shape of the virus particles. The same thresholds and parameters were used for all images.

#### Minigenome reporter assay

Cells were transfected with pCAGGS expression plasmids encoding PB2, PB1, PA, and NP (from PR8, Eng195, Vic75, or Tky05 viruses as indicated in Figure 3 and 4) using Lipofectamine 3000 (Invitrogen). A pPolI-Firefly luciferase viral reporter plasmid and a pCAGGS-Renilla luciferase plasmid, which acts as a transfection efficiency control, were also transfected. Plasmids were transfected in a 2:2:1:4:4:2 (PB2:PB1:PA:NP:Firefly:Renilla) ratio, where 1=10ng, 20ng, 80ng in a 48-well, 24-well, or 6-well plate, respectively. For complementation, pCAGGS plasmids encoding HMGB2 or RUVBL2 (amounts indicated in Figure S5) were also transfected.

For luciferase readout, cells were lysed in passive lysis buffer (Promega) 20hrs post transfection. Bioluminescence was measured using a dual-luciferase assay kit (Promega) on a FLUOstar Omega plate reader (BMG LabTech). Firefly luciferase activity was normalized to Renilla luciferase activity. For RT-qPCR readout, described below, cells were lysed in buffer RLT (Qiagen) supplemented with 10% BME. For immunoblot analysis, described below, cells were lysed with homemade radioimmunoprecipitation assay (RIPA) buffer (50mM Tris (pH 7.4), 150 mM NaCl, 1% NP-40, 0.5% sodium deoxycholate, 0.1% sodium dodecyl sulfate) supplemented with EDTA-free protease inhibitor cocktail tablet. Lysates were centrifuged at 13000xg for 10min, and supernatants were mixed with 4xLaemmli sample buffer with 10% BME.

#### Immunoblot

Proteins were separated by SDS-PAGE on mini protean TGX precast gels 4-20% (Bio-Rad) followed by semi-dry transfer onto PVDF membranes (Low fluorescence 0.2µM; Amersham). Primary antibodies used were rabbit IZ-vinculin (EPR8185; Abcam), rabbit IZ-PB2 (GTX125926; Genetex), rabbit IZ-HMGB2 (EPR6301; Abcam), rabbit IZ-RUVBL2 (10195-1-AP; ThermoFisher Scientific), mouse IZ-tubulin (AB7291; Abcam), mouse IZ-FLAG (F1804, Sigma). Secondary antibodies were IRDye 680RD goat anti-mouse (IgG) (AB175775; Abcam) and IRDye 800CW goat anti-rabbit (IgG) (926-32211; Li-Cor Biosciences). Membranes were visualised using an Odyssey imaging system (LiCor Biosciences).

#### RT-qPCR

Following cell lysis in buffer RLT with 10% BME, total RNA was extracted using a RNeasy RNA extraction kit (Qiagen) with 30min DNase I digestion (Qiagen). A549 cells were homogenized with QIAshredder columns (Qiagen) prior to RNA extraction.

Segment 4 (infection setup) or Firefly (minigenome setup) vRNA, cRNA, and mRNA accumulation was quantified using a tagged RT-qPCR method, developed by Kawakami et al. (65). vRNA, cRNA, mRNA, and GAPDH (internal control) cDNA was synthesised using RevertAid H Minus Reverse Transcriptase (ThermoFisher Scientific) according to the manufacturer’s instructions. For each reaction, 200ng RNA and tagged primers targeting vRNA or cRNA, tagged poly(dT) (mRNA), or untagged poly(dT) (GAPDH) was used. Tagged cDNA was then quantified in triplicate using Fast SYBR green master mix (ThermoFisher Scientific) on a Viia 7 real-time PCR system. Primers used for cDNA and qPCR reactions are shown in Table S4. The 2^-ΔΔCT^ method with GAPDH expression as internal control was used to calculate the fold change in vRNA, cRNA, and mRNA accumulation relative to input (0hpi) (infection setup) or-PB2 control (minigenome setup).

#### cRNP stabilization assay

A549 WT, HMGB2-KO, and RPAP2-KO cells were cultured in 24-well plates. Cells were transfected using Lipofectamine 3000 with pCAGGS expression plasmids encoding PR8 PB2, PB1 D446Y (catalytically dead), PA, and NP in a 2:2:1:4 (PB2:PB1:PA:NP) ratio, where 1=20ng in a 24-well plate. 20hrs post transfection, cells were preincubated in media containing cycloheximide (CHX, 100µg/mL; Sigma) for 1hr. Synchronised infections were performed, as described above, with addition of CHX to media used for preincubation, viral inoculums, and subsequent prewarmed media.

#### Inflachromene treatment

Maximum inflachromene (ICM; Cayman chemical) concentration (0-40µM) tolerated by eHAP WT cells was determined using an MTS cell proliferation kit (Abcam), according to the manufacturer’s instructions. For multi-cycle infections, ICM (20µM) or DMSO was added to viral inoculums, and subsequent prewarmed SF media. At the appropriate timepoint, supernatants were harvested, and infectious titres determined by plaque assay on MDCK cells. For synchronised single-cycle infections, ICM (20µM) or DMSO was added to media used for preincubation, viral inoculums, and subsequent prewarmed media. For minigenome reporter assays, eHAP WT cells were treated with ICM or DMSO immediately prior to plasmid transfection (concentrations indicated in Figure 5).

### Quantification and statistical analysis

#### Statistical analysis

Statistical analysis was performed using GraphPad Prism (v10.2.1) using an unpaired *t* test or multiple unpaired *t* tests, corrected for multiple comparisons using the FDR, as indicated in figure legends. Differences were considered significant at p-values at or below 0.05, as indicated in figure legends. Error bars indicate SD from triplicate experiments, unless otherwise stated in figure legends. Further information about n numbers and statistical tests is presented in figure legends.

## Notes

### Competing Interest Statement

The authors have declared no competing interest.

