## Supplementary Figures 1-6 for "Influenza A virus polymerase co-opts distinct sets of host proteins for RNA transcription or replication"

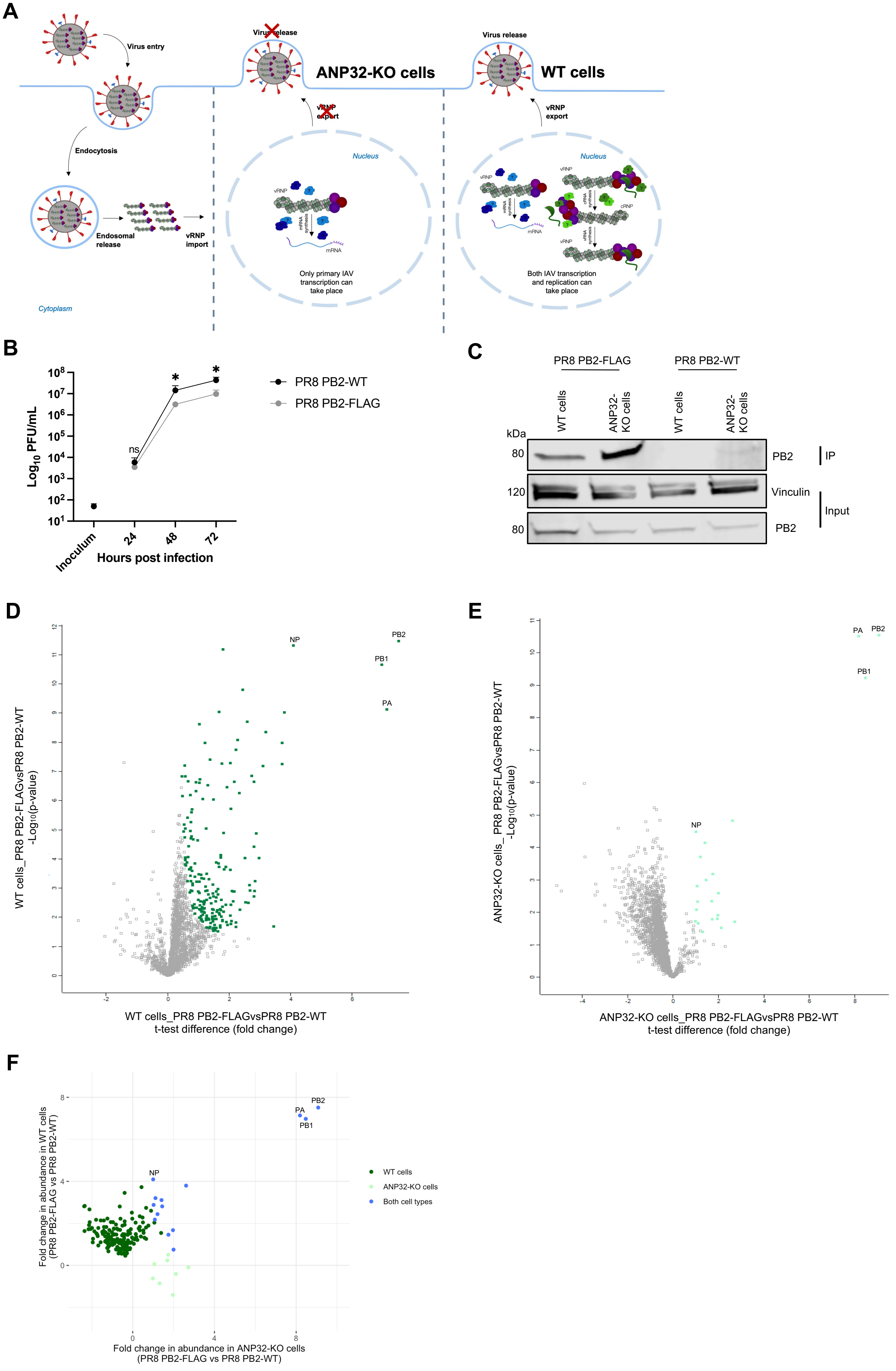
A
Virus entry
Endocytosis
Endosomal release
vRNP import
Cytoplasm
Virus release
ANP32-KO cells
vRNP export
Nucleus
vRNP
mRNA synthesis
mRNA
Only primary IAV transcription can take place
Virus release
WT cells
vRNP export
Nucleus
vRNP
cRNA synthesis
cRNP
vRNA synthesis
Both IAV transcription and replication can take place
B
Log10 PFU/mL
ns
*
*
PR8 PB2-WT
PR8 PB2-FLAG
Inoculum
24
48
72
Hours post infection
C
PR8 PB2-FLAG
PR8 PB2-WT
WT cells
ANP32-KO cells
WT cells
ANP32-KO cells
kDa
80
120
80
PB2
IP
Vinculin
Input
PB2
D
WT cells_PR8 PB2-FLAGvsPR8 PB2-WT
-Log10(p-value)
NP
PB2
PB1
PA
WT cells_PR8 PB2-FLAGvsPR8 PB2-WT
t-test difference (fold change)
E
ANP32-KO cells_ PR8 PB2-FLAGvsPR8 PB2-WT
-Log10(p-value)
PA
PB2
PB1
NP
ANP32-KO cells_PR8 PB2-FLAGvsPR8 PB2-WT
t-test difference (fold change)
F
Fold change in abundance in WT cells
(PR8 PB2-FLAG vs PR8 PB2-WT)
PB2
PA
PB1
NP
WT cells
ANP32-KO cells
Both cell types
Fold change in abundance in ANP32-KO cells
(PR8 PB2-FLAG vs PR8 PB2-WT)

**Figure S1. Identification of host proteins enriched in WT cells, ANP32-KO cells, or both WT and ANP32-KO cells.**

**A)** Schematic of the IAV lifecycle in eHAP WT and ANP32-KO cells. In ANP32-KO cells, the IAV lifecycle is restricted and only steps upstream of and including primary transcription can occur.

**B)** Virus replication in eHAP WT cells infected with PR8 PB2-WT or PR8 PB2-FLAG (MOI=0.001). Supernatants were harvested at indicated timepoints post infection, and titres were determined by plaque assay on MDCK cells. Data is shown as mean ± SD of n=3 technical repeats, representative of n=3 biological repeats. Statistical significance was assessed at each timepoint using an unpaired *t* test. ns, not significant; *, p<0.05.

**C)** Western blot of inputs and immunoprecipitations (IPs) from WT and ANP32-KO cells infected with PR8 PB2-FLAG or PR8 PB2-WT (MOI=5). Three times more ANP32-KO cell lysate than WT cell lysate was used. Cells were lysed at 3hpi. Expression of PB2 and vinculin (cellular control) was detected.

**D and E)** Volcano plots showing proteins co-precipitating from PR8 PB2-FLAG-infected vs PR8 PB2-WT-infected WT cells (D) and ANP32-KO cells (E). Proteins enriched in PR8 PB2-FLAG-infected cells ($S_0$=0.2, FDR=0.05) are shown in green (D) and light green (E).

**F)** Fold change in abundance of proteins above background in WT cells (proteins in greens in (D)) and ANP32-KO cells (proteins in light green in (E)). Proteins enriched in WT cells only are shown in green, in ANP32-KO cells only are shown in light green, and in both WT and ANP32-KO cells in blue.

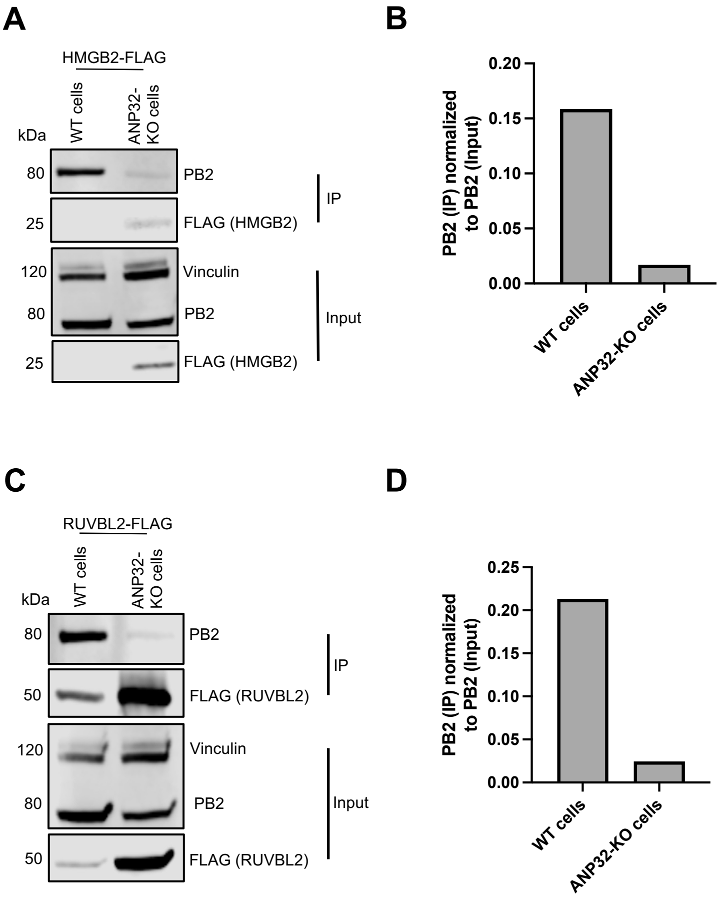


**Figure S2. PB2 co-precipitates with HMGB2 and RUVBL2 to a greater extent in WT cells than ANP32-KO cells.**

**A and C)** Western blot of inputs and immunoprecipitations (IPs) from eHAP WT and ANP32-KO cells transfected with HMGB2-FLAG (A), RUVBL2-FLAG (C) and infected with WT PR8 (MOI=5). Three times more ANP32-KO cell lysate than WT cell lysate was used. Cells were lysed at 3hpi. Expression of PB2, FLAG (HMGB2/RUVBL2), and vinculin (cellular control) was detected.

**B)** Quantification of the amount of PB2 in the IP normalized to the amount of PB2 in the input in (A).

**D)** Quantification of the amount of PB2 in the IP normalized to the amount of PB2 in the input in (C).

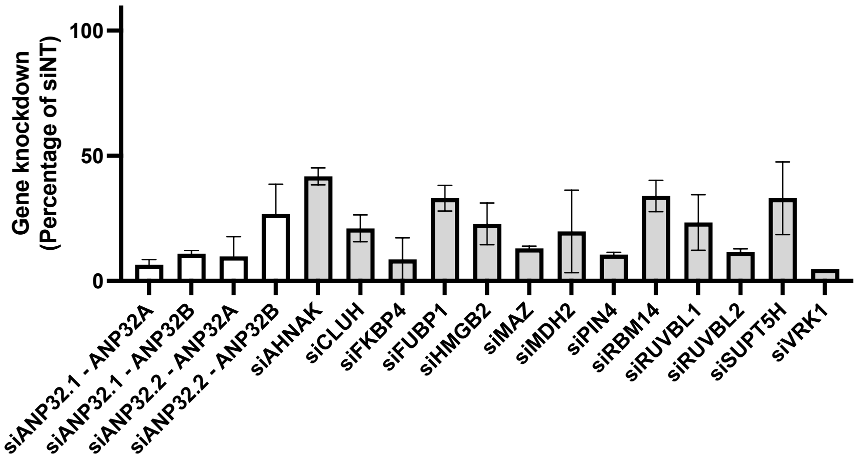


**Figure S3. siRNAs targeting selected genes result in gene knockdown at the mRNA level.** mRNA levels of target genes following siRNA transfection. Data is shown as the percentage gene knockdown relative to mRNA levels in cells transfected with a non-targeting siRNA (siNT). Data is shown as mean ± SD of n=3 technical repeats.

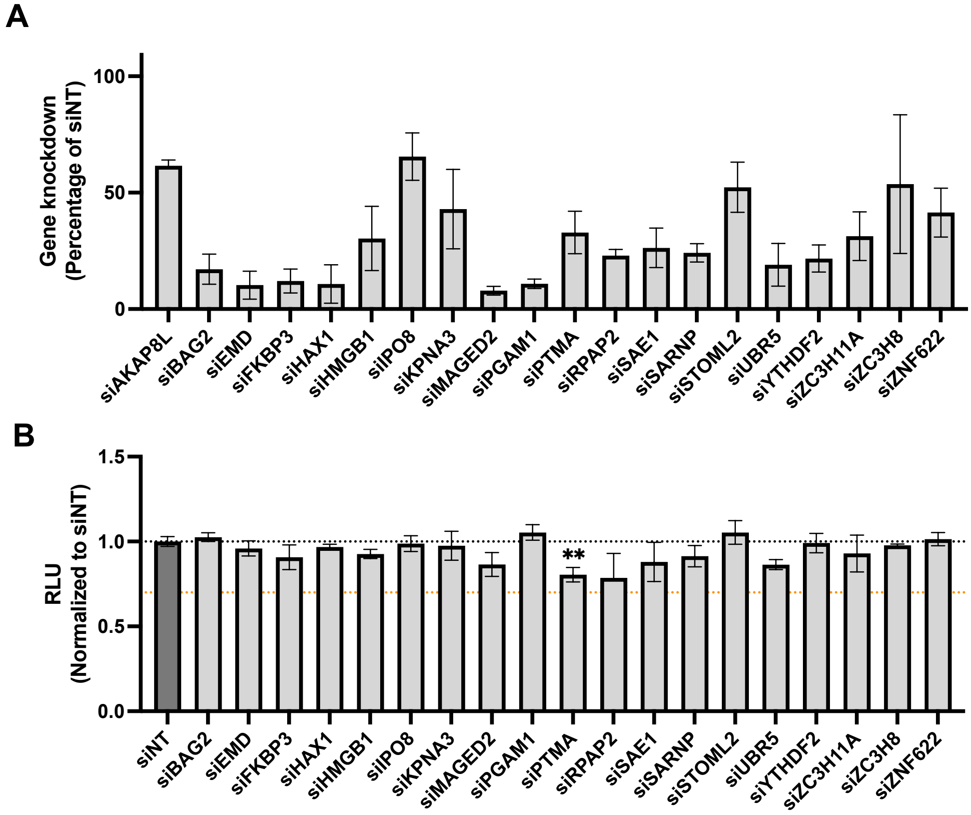


**Figure S4. siRNAs targeting selected genes result in gene knockdown at the mRNA level without affecting cell viability.**

**A)** mRNA levels of target genes following siRNA transfection. Data is shown as the percentage gene knockdown relative to mRNA levels in cells transfected with a non-targeting siRNA (siNT). Data is shown as mean ± SD of n=3 technical repeats.

**B)** Cell viability determined using a CellTiter-Glo assay and shown as luciferase activity (RLU) normalized to siNT. Black dotted line indicates cell viability of siNT-treated cells. Orange dotted line indicates 70% cell viability threshold. Data is shown as mean ± SD of n=3 technical repeats, representative of n=3 biological repeats. Statistical significance was assessed compared to siNT using an unpaired *t* test. ns, not significant; **, $p<0.01$.

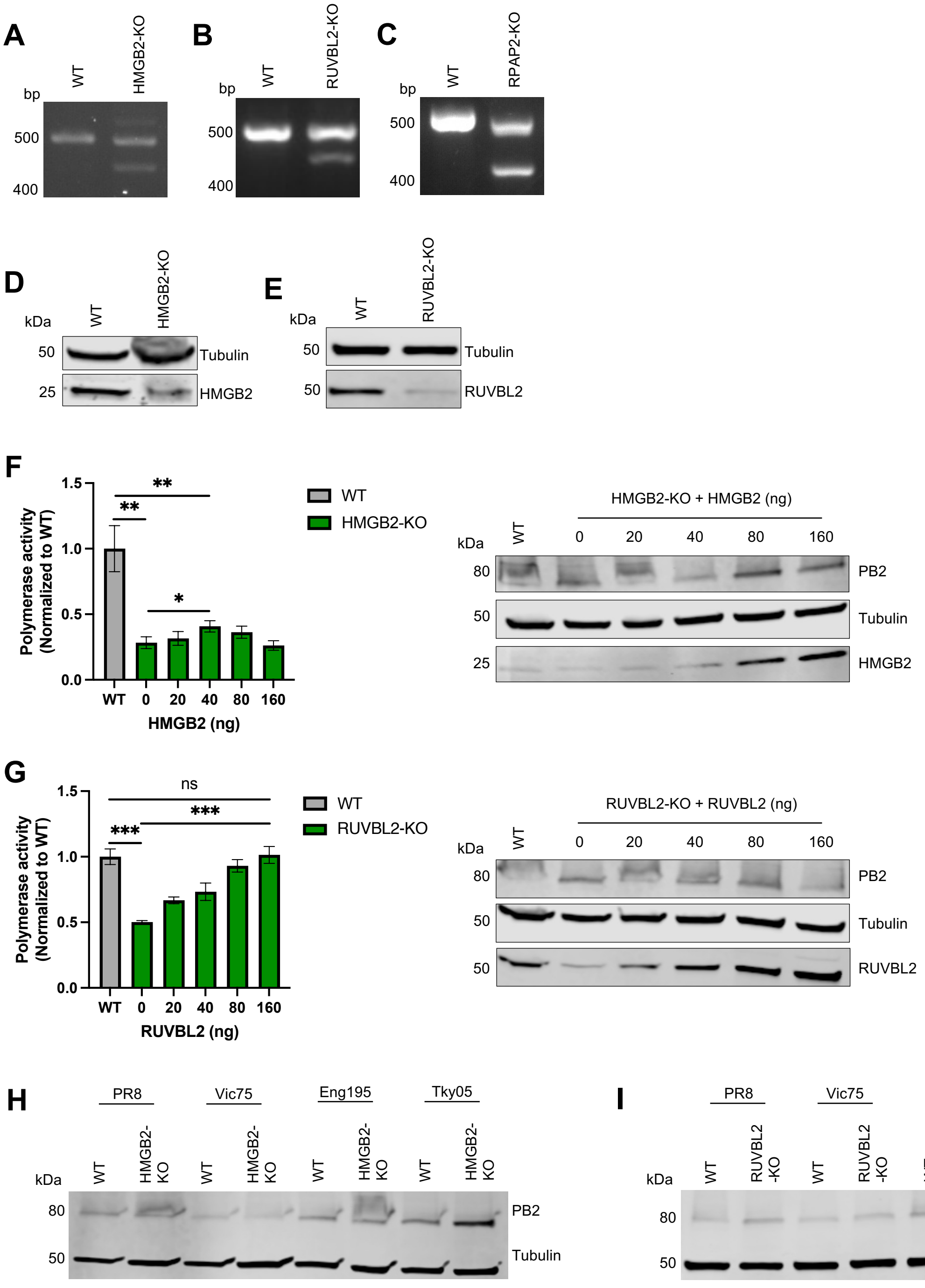


**Figure S5. Polymerase activity is reduced in HMGB2-KO and RUVBL2-KO cells.**

**A-C)** Validation of HMGB2-KO, RUVBL2-KO, and RPAP2-KO single-cell clones. PCR products amplified from genomic DNA from WT, HMGB2-KO (A), RUVBL2-KO (B), and RPAP2-KO (C) cells.

**D and E)** Validation of HMGB2-KO and RUVBL2-KO single-cell clones by western blot. Expression of HMGB2 (D), RUVBL2 (E), and tubulin (cellular control) was detected.

**F and G)** Complementation with HMGB2 and RUVBL2 partially or fully restores polymerase activity to WT levels, respectively. Minigenome reporter assay in A549 WT and HMGB2-KO cells (F) or A549 WT and RUVBL2-KO cells (G). Cells were transfected with plasmids to reconstitute PR8 polymerase activity and plasmids expressing HMGB2 (F) and RUVBL2 (G) (amounts indicated in figure). Firefly luciferase activity was normalized to co-transfected Renilla luciferase levels. Accompanying western blots show HMGB2 (F), RUVBL2 (G), PB2, and tubulin (cellular control) protein expression for each condition. Data is shown as mean ± SD of n=3 technical repeats, representative of n=3 biological repeats. Statistical significance was assessed compared to WT using an unpaired *t* test. ns, not significant; *, p<0.05; **, p<0.01; ***, p<0.001.

**H and I)** Western blots accompanying minigenome reporter assay in Figure 3B-I, showing PR8, Vic75, Eng195, and Tky05 PB2 expression as well as tubulin (cellular control) in WT and HMGB2-KO cells (H) or WT and RUVBL2-KO cells (I).

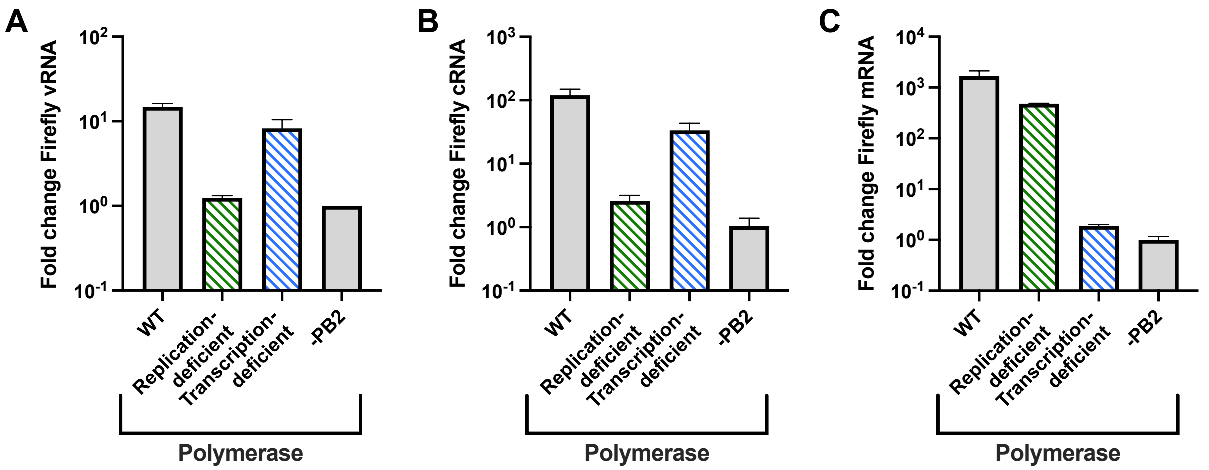


**Figure S6. Characterization of replication-deficient and transcription-deficient polymerase mutants.**
**A-C)** Accumulation of Firefly luciferase vRNA (A), cRNA (B), and mRNA (C) in eHAP WT cells. Cells were transfected with plasmids to reconstitute WT PR8 vRNPs, replication-deficient vRNPs (PA E410A) (green striped), and transcription-deficient vRNPs (PA D108A) (blue striped). vRNA template provided was a pPolI-Firefly luciferase plasmid. Empty plasmid was substituted for PB2 as a negative control (-PB2 condition). Data is shown as the fold change over -PB2 condition. Data is shown as mean ± SD of n=3 technical repeats, representative of n=3 biological repeats.
