## Supplementary Table 2 for "Influenza A virus polymerase co-opts distinct sets of host proteins for RNA transcription or replication"

**Table S2. Prioritisation of host proteins for onwards validation.** The host function, localisation, and role in the IAV lifecycle of host proteins enriched specifically in WT cells, in both WT and ANP32-KO cells, or specifically in ANP32-KO cells**.** Proteins in bold were previously identified in IAV screens (proteomics or functional) or previously shown to play a role in the IAV lifecycle. Proteins in green were selected for onwards validation as replication co-factors. Proteins in blue were selected for onwards validation as transcription co-factors. Viral proteins are not shown. Related to Figure 1.

| **Cell type host protein was identified in** | **GO term** | **P-value** | **Protein names** | **Role in IAV lifecycle** |
| --- | --- | --- | --- | --- |
| Host proteins enriched specifically in WT cells | **Biological process:** ATP metabolic process  GO:0046034 | 2.50E-04 | NDUFA9, NDUFV2, UQCRC1, PGAM1, STOML2, HK1, PGK1, LDHA | N/A |
|  | **Biological process:** Regulation of protein ubiquitination  GO:0031396 | 0.0281 | **UBR5**, UBQLN1, UBE2N, **PDCD6** | UBR5 identified as PA and PB2 interactor (1-4). PDCD6 identified as PB2 interactor (3). UBR5 and PDCD6 found to be proviral regulators of IAV lifecycle (1, 5). |
|  | **Biological process:** Regulation of translation initiation  GO:0006446 | 0.0224 | **YTHDF2**, EIF4H, EIF4G2 | YTHDF2 identified as PB2 interactor and proviral factor in IAV lifecycle (2, 6). |
|  | **Biological process:** RIG-I signalling pathway  GO:0039529 | 0.0139 | **PHB1**, PHB2, **CLPB** | PHB1 shown to have a proviral role in IAV lifecycle (4). CLPB identified as a PB2 interactor (3). |
|  | **Biological process:** Protein folding  GO:0006457 | 1.11E-08 | **HSPA9**, **HSPA8**, HSPE1, **HSP90AA1**, **HSPD1**, **HSPA4**, **HSPB1**, **HSPA1B**, **DNAJA1**, **DNAJA2**, **DNAJA3**, **DNAJB11**, **BAG2**, FKBP4, CLPX, PDIA3, **P4HB**, CALR | Proteins in bold identified as polymerase interactors (1-4, 7) or as regulators of the IAV lifecycle (8-11). HSP90AA and DNAJA1 act as polymerase chaperones (10, 11). |
|  | **Biological process:** Transcription  GO:0006351 | 0.0025 | **POLR2B**, **POLR2C**, **POLR2D**, **POLR2E**, **POLR2G**, **POLR2H**, **POLR2L**, **SUPT5H**, TRIP13, ZC3H8, MAZ, **PTMA** | Proteins in bold identified as polymerase interactors (1-4, 7) or as regulators of the IAV lifecycle (8, 9, 12). Host RNAPII is important for viral transcription (13). |
|  | **Cellular component:** Nuclear envelope  GO:0005635 | 0.0227 | IPO8, **DHRS2**, **KPNA3**, CACYBP, **LBR**, **EMD**, RTN4, CLIC1, **HAX1**, RB1CC1 | Proteins in bold shown to be polymerase interactors (2, 3, 7) or regulators of IAV lifecycle (8, 14). |
|  | **Cellular component:** Nucleolus  GO:0005730 | 4.30E-04 | NOL6, STAU2, **MAGED2**, ZNF622, PIN4, **AKAP8**, CDKN2AIPNL, HJURP, **RBM14**, VRK1, PHF8, **SNRPB2, TIMM44, TP53,** ZNF622, **SNRPB2** | Proteins in bold identified as polymerase (2, 4, 7). RBM14 shown to have a proviral role in IAV polymerase activity (15). TIMM44 and TP53 shown to be regulators of the IAV lifecycle (8, 12). |
|  | **Cellular component:** Prohibitin complex  GO:0035632 | 0.0143 | **PHB1**, PHB2 | PHB1 shown to have a proviral role in IAV lifecycle (4). |
|  | **Cellular component:** Proteasome core complex  GO:0005839 | 8.05E-06 | **PSMA1**, PSMA2, PSMA3, PSMB1, PSMB5, PSMB4 | PSMA1 identified as PB2 interactor and proviral factor in IAV lifecycle (4, 8). |
|  | **Cellular component:** R2TP complex  GO:0097255 | 0.0438 | **RUVBL1**, **RUVBL2** | Identified as interactors of PB2, PB1, and NP (1, 7, 12, 16, 17). Shown to be regulators of IAV lifecycle (1, 16). |
|  | **Cellular component:** RNA polymerase II, holoenzyme  GO:0016591 | 4.09E-07 | **POLR2B**, **POLR2C**, **POLR2D**, **POLR2E**, **POLR2G**, **POLR2H**, **POLR2L** | Proteins in bold identified as polymerase interactors (1-4, 7) or as regulators of the IAV lifecycle (8). |
|  | Other |  | SEC63, HISTH2B, SPC25, SSR1, PRKCSH, RALA, SARS2, GPC4, **CLUH**, TLN1, HADHB, ECHS1, PYCR1, RAD23B, CAP1, SLC16A1, **CSTF3**, NME1, NSDHL, GTPBP6, ME2, **MLF2**, SDF4, **SAE1**, **PPM1G**, ZC3H11A, HARS, ESD, NES, PDS5A, GPN3, CNN2, **AIFM1**, PHKG2, CALM, PFN1, CS, MDH2, ACO2, MYEF2, PEBP1, ATIC, **ALDH18A1**, OAT, RCN1, RAB2, SMS, SNX2, SARNP, PSAT1, TECR, THAP11, **ALDH1B1**, PMPCB, **MAP1B**, PDHB, KIF23, SNRPG, **TUFM**, PAPSS1, **AHNAK**, FKBP3, CNDP2, SF1, AP4S1, ERAL1, **CALU**, SEPTIN-9, BLVRA, ACTA1, SMARCE1, RAB1, HDGF, ANXA5, TFRC, **GLUD1**, C4, ETFA, **CYCS**, FUBP1, ENAH, TALDO1, MACF1, AURKA, GSTP1, AMD1, **HMGB1, HMGB2** | Proteins in bold shown to be polymerase interactors (1-4, 7, 18) or regulators of IAV lifecycle (8, 9, 12, 18). |
| Host proteins enriched in both WT and ANP32-KO cells | **Biological process:** Transcription  GO:0006351 | 0.0025 | **POLR2A**, **POLR2I**, **POLR2J**, RPAP2 | Proteins in bold identified as polymerase interactors (7). Host RNAPII is important for viral transcription (13). |
|  | **Cellular component:** RNA polymerase II, holoenzyme  GO:0016591 | 4.09E-07 | **POLR2A, POLR2I, POLR2J,** RPAP2 | Proteins in bold identified as polymerase interactors (7). Host RNAPII is important for viral transcription (13). |
|  | Other |  | **SEC61B, PJA2**, **MAGED1**, **RNF219**, **AKAP8L**, **HSBP1** | Proteins in bold shown to be polymerase interactors (1-4). |
| Host proteins enriched specifically in ANP32-KO cells | **Biological process:** ATP metabolic process  GO:0046034 | 2.50E-04 | NDUFAB1 | N/A |
|  | **Cellular component:** Nucleolus  GO:0005730 | 4.30E-04 | CSTB, GOLGA3 | N/A |
|  | Other |  | **HIST1H2A**, MYO5B, MYL1, SNRPG, RPL26L1 | HIST1H2A shown to be regulator of IAV infection (12). |
